# FilAdh: degradation-aware analysis of focal adhesions and actin stress fibers

**DOI:** 10.64898/2026.09.28.755011

**Authors:** Guillermo Rey-Paniagua, Carlos García-López-de-Haro, María Hernández-Martínez, Dariusz Lachowski, Arrate Muñoz-Barrutia

## Abstract

Focal adhesions (FAs) and actin stress fibers (SFs) are mechanically coupled but often analyzed separately. We present FilAdh, a unified framework for FA instance segmentation, SF instance reconstruction, morphometry, and FA-anchorage-based SF classification. Across 57 held-out HeLa, SUIT-2, and MRC-5 cell line images, FilAdh achieved the highest FA Dice, intersection over union (IoU), and recall and the highest SF clDice and skeleton precision among benchmarked methods; paired *t* tests with Holm correction indicated significant gains over all compared methods for FA Dice, IoU, and recall, and over classical methods for SF centerline metrics, with no significant differences detected relative to the deep-learning comparators. Conditional on expert-curated FA and SF masks, SFs were assigned to dorsal, ventral, and peripheral classes with 81.5%–87.3% accuracy. Controlled Poisson-Gaussian noise, Gaussian defocus, and point-spread-function blur revealed distinct, structure-specific errors in overlap and object-level measurements. By linking segmentation and reconstruction to count, area, length, apparent width, and structural class, FilAdh offers an interpretable view of the FA-actin structural system and makes degradation-dependent measurement biases explicit.

**BIGGER PICTURE:** Cells sense and respond to their physical environment through focal adhesions and actin stress fibers. The two form a mechanically coupled system, yet they are usually analyzed separately, with workflows re-tuned for each new set of images. FilAdh analyzes both within a single framework, tailoring segmentation, reconstruction, and classification to each structure’s geometry: compact adhesion patches versus thin, extended fiber networks. Testing on expert-annotated references and under controlled degradations that mimic low signal, focus errors, and optical blur shows that these common imaging imperfections distort adhesion and fiber measurements in different ways. The same design principle—matching the analysis to each structure’s shape and judging methods by how errors propagate into the final biological measurements—may benefit other bioimage-analysis problems. Released code, trained weights, and fixed data splits make all comparisons reproducible and extensible.

**HIGHLIGHTS:**

- FilAdh jointly quantifies focal adhesions and actin stress fibers
- FilAdh achieves the highest benchmarked FA overlap and comparable SF topology
- Matched synthetic degradations reveal geometry-specific measurement errors
- FA-SF spatial context enables dorsal, ventral, and peripheral SF classification

## INTRODUCTION

Focal adhesions (FAs) are integrin-based protein assemblies that couple the extracellular matrix to the actin cytoskeleton. They act as mechanical anchors and signaling hubs, allowing cells to transmit actomyosin-generated forces, sense matrix mechanics, and coordinate spreading and migration. Their molecular organization spans integrins, force-transmission proteins such as talin and vinculin, and actin-associated layers; adhesion growth and maturation are therefore tightly coupled to force generation and stress-fiber organization^1–5^.

Actin stress fibers (SFs) are prominent actin filament bundles that organize cellular tension and force transmission, with distinct architectures and anchorage patterns. In the 2D operational annotation scheme used here (Figure 1), dorsal SFs contact a focal adhesion at one end and are oriented approximately perpendicular to the nearest apparent cell membrane. Ventral SFs contact focal adhesions at both ends, lie in the central region of the cell, and are oriented approximately tangent to the nearest apparent membrane. Peripheral SFs lie at the cell periphery and contact focal adhesions at both ends. These arrangements support different modes of force transmission, cell-shape control, and migration^6–8^. Joint FA–SF analysis can therefore reveal structural relationships that are not accessible when each structure is measured separately.

**Figure 1:**
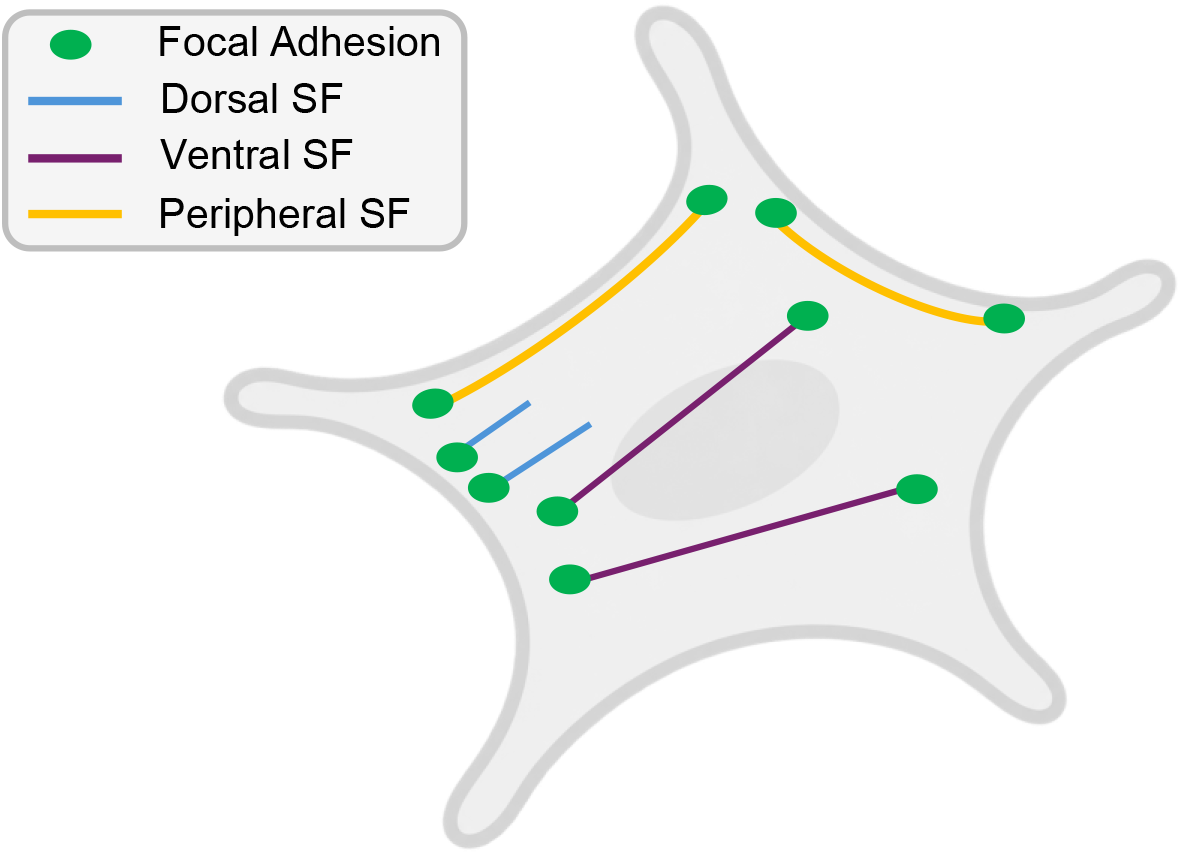
Operational two-dimensional classification of actin stress fibers. Schematic representation of the dorsal, ventral, and peripheral SF classes used in this study, defined by focal-adhesion anchorage, position within the cell, and orientation relative to the apparent cell boundary.

The FA–SF interface is also relevant to mechanically regulated disease processes. Changes in extracellular-matrix stiffness and actomyosin regulation alter adhesion signaling, cellular contractility, and invasive behavior^9,10^. Quantitative FA–SF phenotypes can thus provide structural readouts for experiments that examine how cells respond to biochemical and mechanical perturbations.

Reliable measurement is difficult for both biological and optical reasons. On the biological side, FAs range from nascent adhesions to mature, elongated focal adhesions and often lie close together, whereas SFs form thin, overlapping networks whose apparent continuity can be interrupted when filament segments leave the focal plane. On the optical side, fluorescence microscopy imposes a trade-off among exposure, temporal resolution, and signal-to-noise ratio^11^. Structural visibility is further reduced by photon noise and photobleaching, by nonuniform background, and by optical blurring introduced by defocus and the point-spread function. Reference annotation introduces additional uncertainty: expert labels may differ, and semi-automated masks inherit preprocessing and thresholding choices^12^.

Existing methods address different parts of this problem. For FAs, the Focal Adhesion Analysis Server (FAAS)^13^ and the Horzum workflow^14^ preprocess images by filtering and contrast enhancement, binarize them by thresholding, and extract FA instances through connected-component analysis. Here, the Horzum workflow is benchmarked through the containerized implementation of Hung et al.^15^, which fixes all parameters before test-set evaluation and thus ensures reproducible execution.

Deep-learning approaches include star-convex instance segmentation and U-Net-based semantic segmentation. These methods reduce manual parameter tuning but retain distinct geometric and post-processing assumptions^16–19^. Table 1 summarizes the FA comparison set.

**Table 1:** FA-method taxonomy and benchmark role. The image-adjusted Horzum work-flow supported reference construction; its fixed-parameter Hung implementation was evaluated quantitatively.

| Family | Method | Core approach | Benchmark use |
| --- | --- | --- | --- |
| **Classical** | FAAS, Berginski et al. (2013)<sup>13</sup> | Filtering, thresholding, and connected-component analysis | User-defined parameters; FA instances and morphometry. |
|  | Horzum et al. (2014)<sup>14</sup> | Background subtraction, local contrast enhancement, Laplacian-of-Gaussian filtering, and thresholding | Reference construction with image-specific tuning and expert correction. |
|  | Horzum implementation, Hung et al. (2022)<sup>15</sup> | Containerized implementation of the Horzum workflow | Fixed test parameters; FA instances and morphometry. |
| **Deep Learning** | Weigert et al. (2020)<sup>16</sup> | Star-convex instance segmentation | Supervised instance-segmentation comparator. |
|  | Mohamed et al. (2024)<sup>17,19</sup> | U-Net and U-Net++ semantic segmentation | Supervised semantic-segmentation comparator. |
| **Proposed** | **FilAdh** | NAFNet-GAN enhancement<sup>20</sup>, Graph-Bottleneck Attention U-Net segmentation, and watershed separation | Pooled training; FA instances and morphometry. |

Stress-fiber analysis has progressed from multiscale line detection and rule-based tracing to deep-learning segmentation^21,22^. An earlier study of MRC-5 cells combined focal-adhesion analysis with an earlier FilamentSensor workflow for filament detection and FA–SF cross-correlation^23^. The present benchmark evaluates FilamentSensor 2.0 for modular detection and reconstruction of straight and curved filaments^22^.

SynSeg and the Filamentous Actin Segmentation Tool (FAST) provide deep-learning alternatives, but they were developed using different supervision strategies and target definitions. SynSeg was introduced within a synthetic-data framework, whereas FAST used region-level filament annotations rather than individually traced filament instances^24,25^. We therefore compared the methods on common held-out images, retraining them on shared reference targets where possible, and used metrics designed for thin-structure topology. Table 2 summarizes the SF comparison set.

**Table 2:** SF-method taxonomy and benchmark role. Classical configurations were selected from predefined parameter grids using validation images and frozen before heldout evaluation. SynSeg and FAST were retrained on the shared splits and SF references.

| Family | Method | Core approach | Benchmark use |
| --- | --- | --- | --- |
| **Classical** | Alioscha-Perez et al. (2016)<sup>21</sup> | Cartoon–texture decomposition and multiscale line detection | Validation-selected configuration; filament segments. |
|  | FilamentSensor 2.0, Hauke et al. (2023)<sup>22</sup> | Modular detection and reconstruction of straight and curved filaments | Validation-selected configuration; filament instances and morphometry. |
| **Deep Learning** | Guo et al. (2026), SynSeg<sup>24</sup> | U-Net segmentation developed within a synthetic-data framework | Retrained on shared splits; binary SF mask. |
|  | Aljapur et al. (2026), FAST<sup>25</sup> | Deep encoder–decoder for filamentous-actin segmentation | Retrained on shared splits; binary SF mask. |
| **Proposed** | **FilAdh** | Graph-Bottleneck Attention U-Net with tangent prediction and orientation-guided instance reconstruction | Pooled training; topology, filament instances, and morphometry. |

Here we present FilAdh, an integrated framework that segments FA instances, reconstructs SF instances, and derives object-level morphometry and anchorage-based SF classes. The framework comprises three specialized but connected branches: FA instance segmentation, orientation-aware SF instance reconstruction, and SF anchorage classification (Figure 2). We designed the study around three questions: whether compact adhesions and filamentous networks can be analyzed within one geometry-aware framework, how segmentation and reconstruction errors affect object-level measurements, and how defined acquisition-related degradations affect the two geometries differently. To answer these questions, we benchmark clean-image performance on a fixed held-out test subset and then evaluate the same source images under matched synthetic degradations. We further assess class assignment using expert-curated FA and SF instances and report hardware-matched inference times for the segmentation networks.

**Figure 2:**
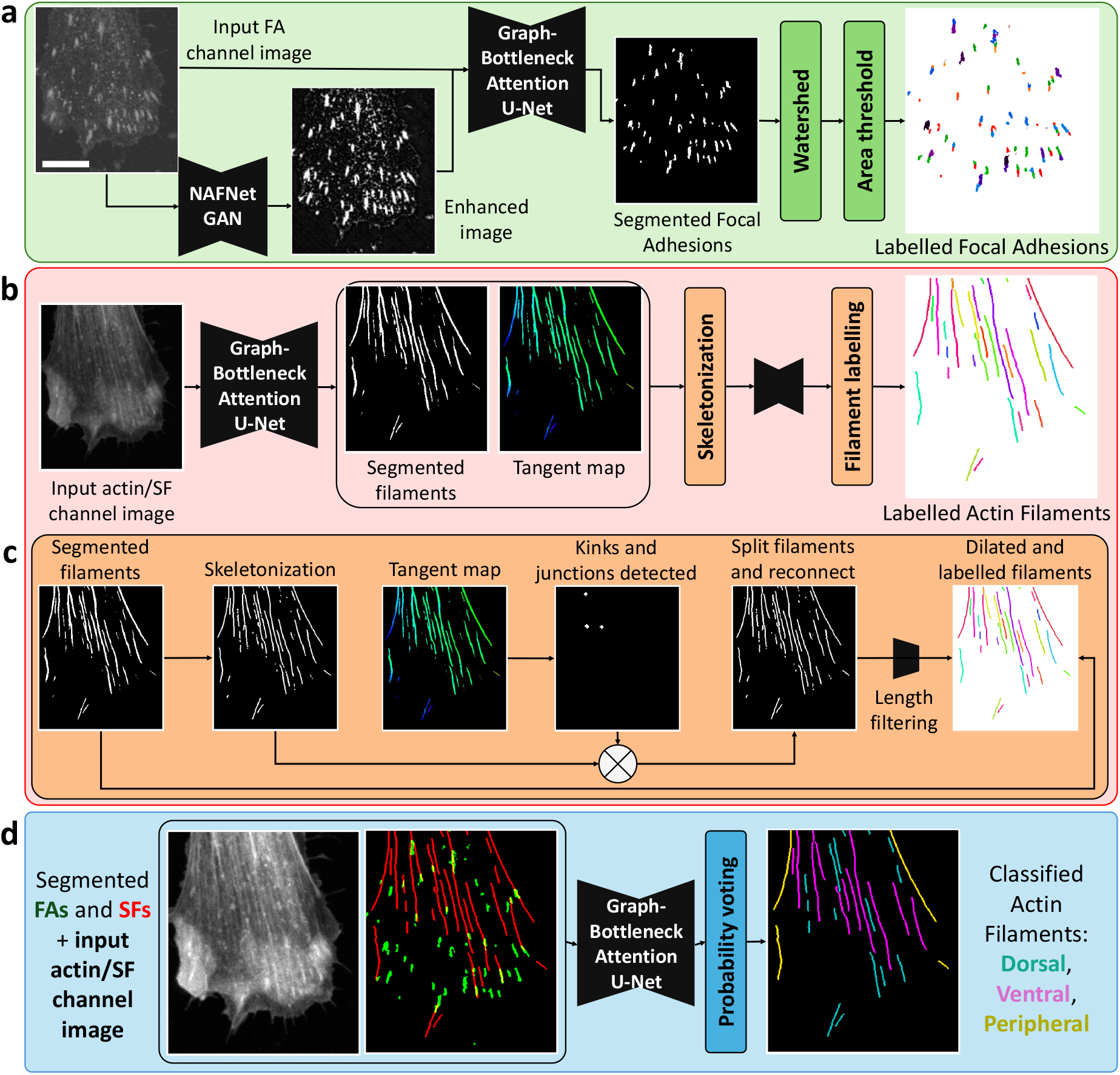
FilAdh integrates FA segmentation, SF reconstruction, and conditional classification. **a**, Raw FA channel images and their NAFNet-GAN-enhanced representations are segmented and separated into FA instances. **b**, The actin/SF channel image is used to predict an SF mask and an undirected tangent map. Cyclic hue encodes the undirected filament orientation *θ* ∈ [0, *π*): red denotes 0^*°*^, blue denotes 60^*°*^, and green denotes 120^*°*^. **c**, Orientation-guided refinement separates paths with incompatible local orientations, reconnects geometrically supported segments, and applies a minimumlength threshold of 1.5 *µ*m to remove short pseudofilaments. **d**, The classifier receives the actin/SF channel image, FA mask, and SF mask, and object-level probability aggregation assigns each filament to the dorsal, ventral, or peripheral class. Display intensities were enhanced for visualization. Scale bar, 10 *µ*m.

## RESULTS

### Heterogeneous microscopy datasets define a common held-out evaluation

Three complementary human cell lines define the evaluation domain. The HeLa images, acquired by spinning-disk confocal microscopy, were obtained from the FAST training and test dataset of Aljapur et al.^25,26^. SUIT-2 pancreatic ductal adenocarcinoma (PDAC) cells were imaged by total internal reflection fluorescence (TIRF) microscopy^27^. MRC-5 human lung fibroblasts were imaged by laser-scanning confocal microscopy using the dataset reported by Hauke et al.^23^.

Across datasets, we refer to the channel reporting focal-adhesion signal as the FA channel and the channel reporting actin stress fibers as the actin/SF channel. Together, the datasets span differences in cell lineage, adhesion density, and filament organization, as well as in acquisition modality, native image dimensions, and signal quality (Figure 3). Dataset size, physical calibration, and the fixed training, validation, and test partitions are summarized in Table 3.

**Table 3:** Cellular systems, microscopy modalities, physical calibration, and fixed data partitions. Image counts are reported as total; training/validation/test.

| Dataset | Cellular system | Imaging modality | Native dimensions (pixels) | Pixel size ($\mu\text{m}/\text{px}$) | Images; split |
| --- | --- | --- | --- | --- | --- |
| HeLa | Cervical carcinoma cells | Spinning-disk confocal microscopy | $1600 \times 1600$ | 0.108 | 297; 214/53/30 |
| SUIT-2 | Pancreatic ductal adenocarcinoma cells | Total internal reflection fluorescence microscopy | $2560 \times 2160$ | 0.108 | 60; 34/9/17 |
| MRC-5 | Human lung fibroblasts | Laser-scanning confocal microscopy | $520 \times 696$ | 0.122 | 48; 30/8/10 |

**Figure 3:**
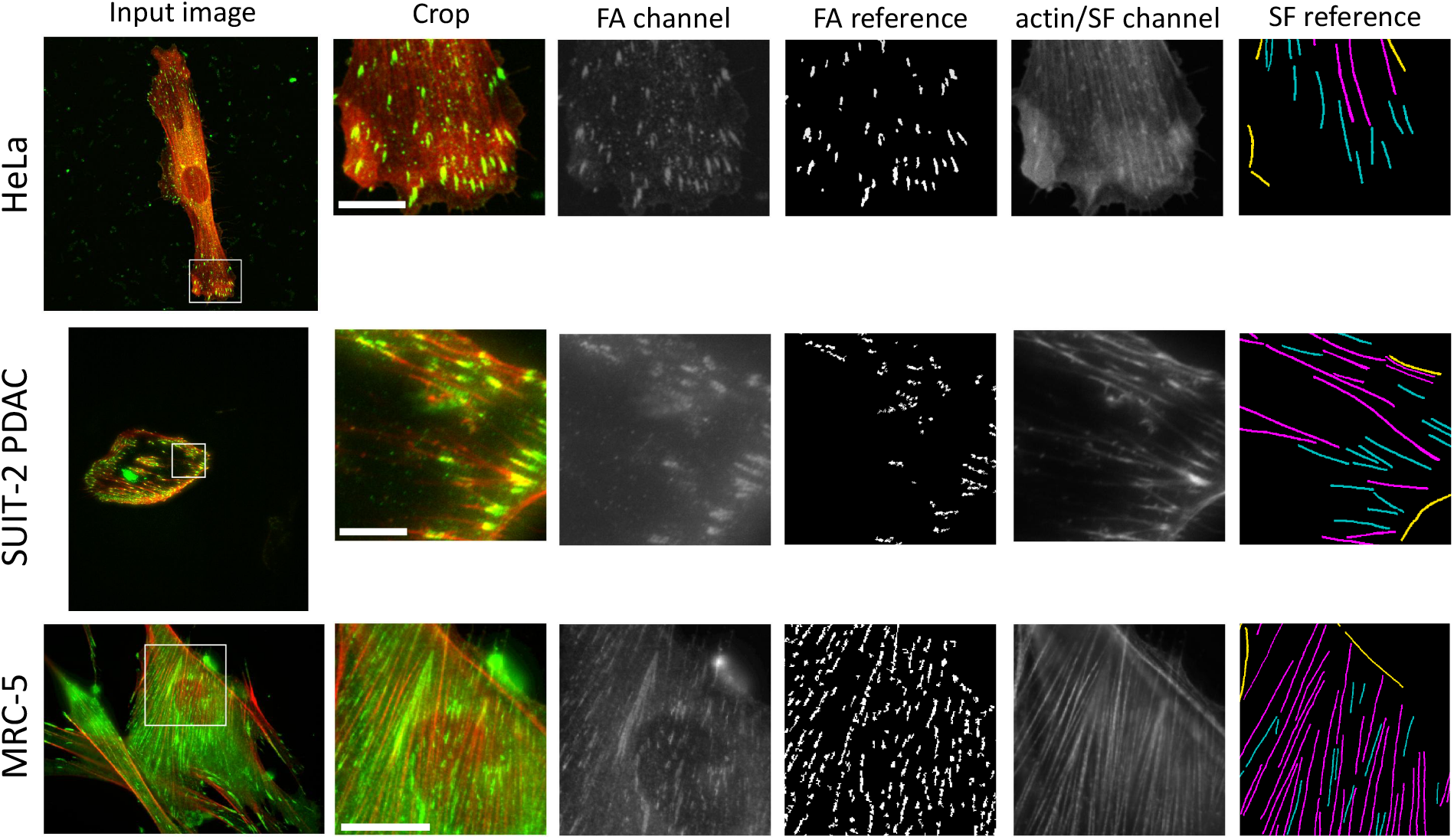
Datasets and expert-curated FA and SF references. Rows show representative HeLa, SUIT-2 PDAC, and MRC-5 images. Columns show the dual-channel field of view, the selected crop, the FA channel, the expert-curated FA reference, the actin/SF channel, and the classified SF reference. In the SF references, teal denotes dorsal fibers, magenta denotes ventral fibers, and yellow denotes peripheral fibers. White boxes indicate the displayed crops. Scale bars, 10 *µ*m.

All reported segmentation, morphometry and runtime comparisons use this fixed heldout test subset: 30 HeLa, 17 SUIT-2 PDAC, and 10 MRC-5 images (57 in total). No training or validation image was included in any reported evaluation.

A single model per task was trained jointly on the pooled data from all three datasets, without dataset-specific fine-tuning. Because the HeLa collection contained substantially more training images (Table 3), SUIT-2 and MRC-5 samples were oversampled by factors of 6 and 7, respectively, within the training and validation loaders to approximately balance dataset sampling. This repetition changed sampling frequency only and did not create additional biological replicates. Each held-out test image contributed one observation to the reported clean-image comparisons.

FA reference masks were generated using an operator-guided workflow adapted from Horzum et al.^14^. The procedure combined sequential filtering and thresholding in Fiji and Python (Table 1), followed by image-specific parameter adjustment and consensus review and correction. SF references were traced manually in Labkit and assigned to dorsal, ventral, or peripheral classes^28^. These classes were assigned according to the operational criteria introduced above and detailed in the STAR Methods. The resulting annotations are treated as expert-curated references rather than parameter-free biological ground truth.

### FilAdh combines enhancement, global feature aggregation, and object reconstruction

Figure 2 summarizes the FilAdh framework. The FA branch enhances the FA channel image, predicts an adhesion mask, and separates touching objects by marker-controlled watershed. The SF branch predicts a filament mask and an undirected local tangent map. This auxiliary orientation field formulation is conceptually related to the flow-based supervision used by Cellpose for instance reconstruction; here, the network instead predicts an undirected local tangent orientation encoded modulo *π* ^29^. The predicted orientations guide the separation of incompatible paths and reconnection of supported segments. The classification branch receives the actin/SF channel image together with FA and SF masks and assigns one structural label to each filament by object-level probability aggregation.

Together, these branches provide FA count and area; SF count, length, and apparent width; and dorsal, ventral, or peripheral classification.

For focal-adhesion enhancement, we reused the previously described NAFNet-GAN architecture and generator objective and retrained the model for the present application^20^.

Raw FA channel images were used as inputs, and the enhanced grayscale representation produced by the image-adjusted Horzum workflow immediately before thresholding was used as the training target (Figure 4)^14^. The discriminator was used during training but not during inference.

**Figure 4:**
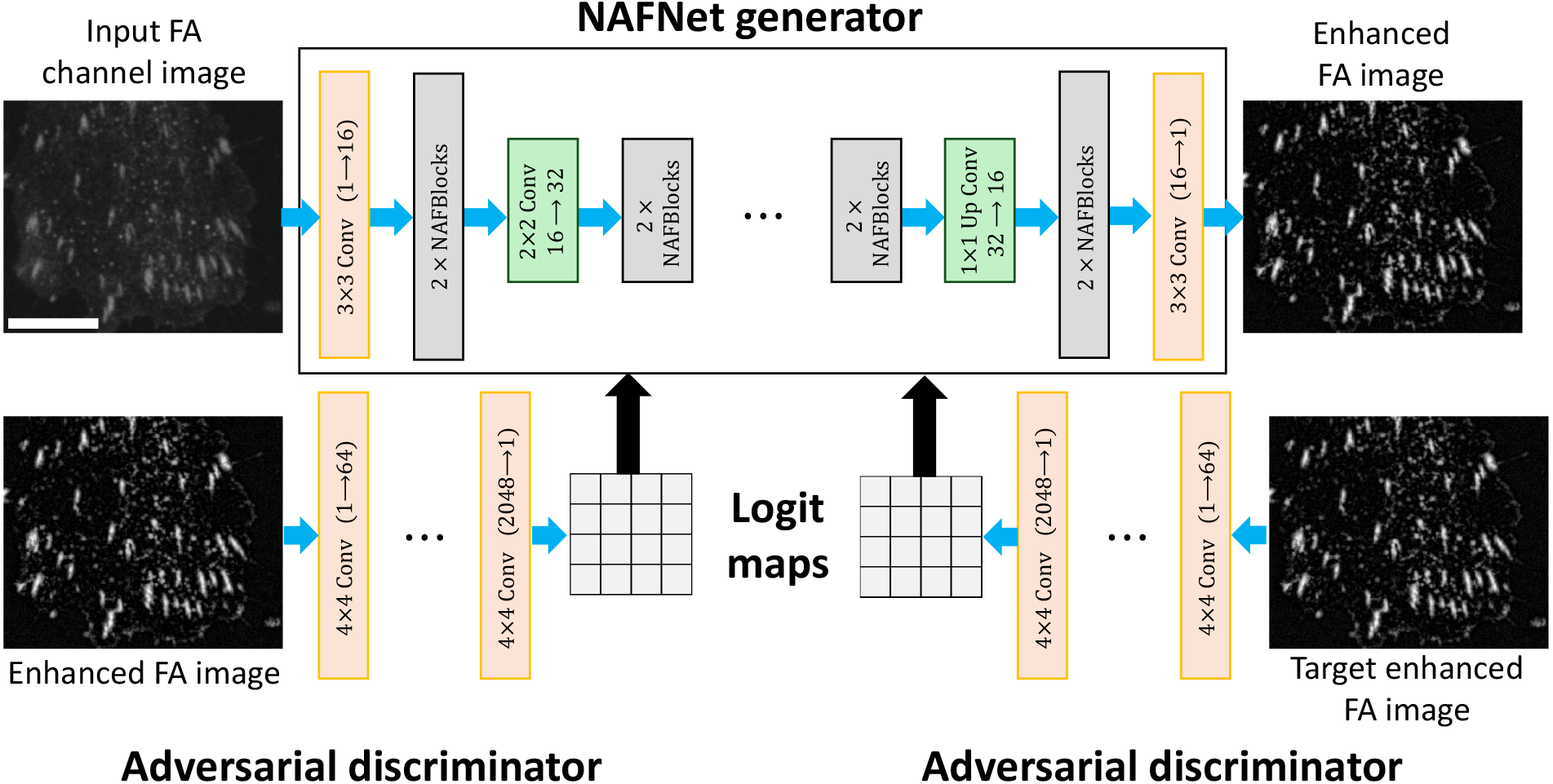
NAFNet-GAN is adapted to a Horzum-derived focal-adhesion enhancement target. The generator maps raw FA channel images to the pre-threshold enhanced representation produced by the image-adjusted Horzum workflow. A shared patch discriminator compares generated and target images during training; only the generator is retained for inference. Scale bars, 10 *µ*m.

The three prediction tasks use compact Graph-Bottleneck Attention U-Net variants (Figure 5). A depthwise-separable encoder extracts local features, a global graph bottleneck aggregates and redistributes contextual information through learned soft spatial-to-node assignments, and attention-gated skip connections restore spatial detail in the decoder. Task-specific heads produce an FA mask, an SF mask with a tangent map, or filament-class probability maps.

**Figure 5:**
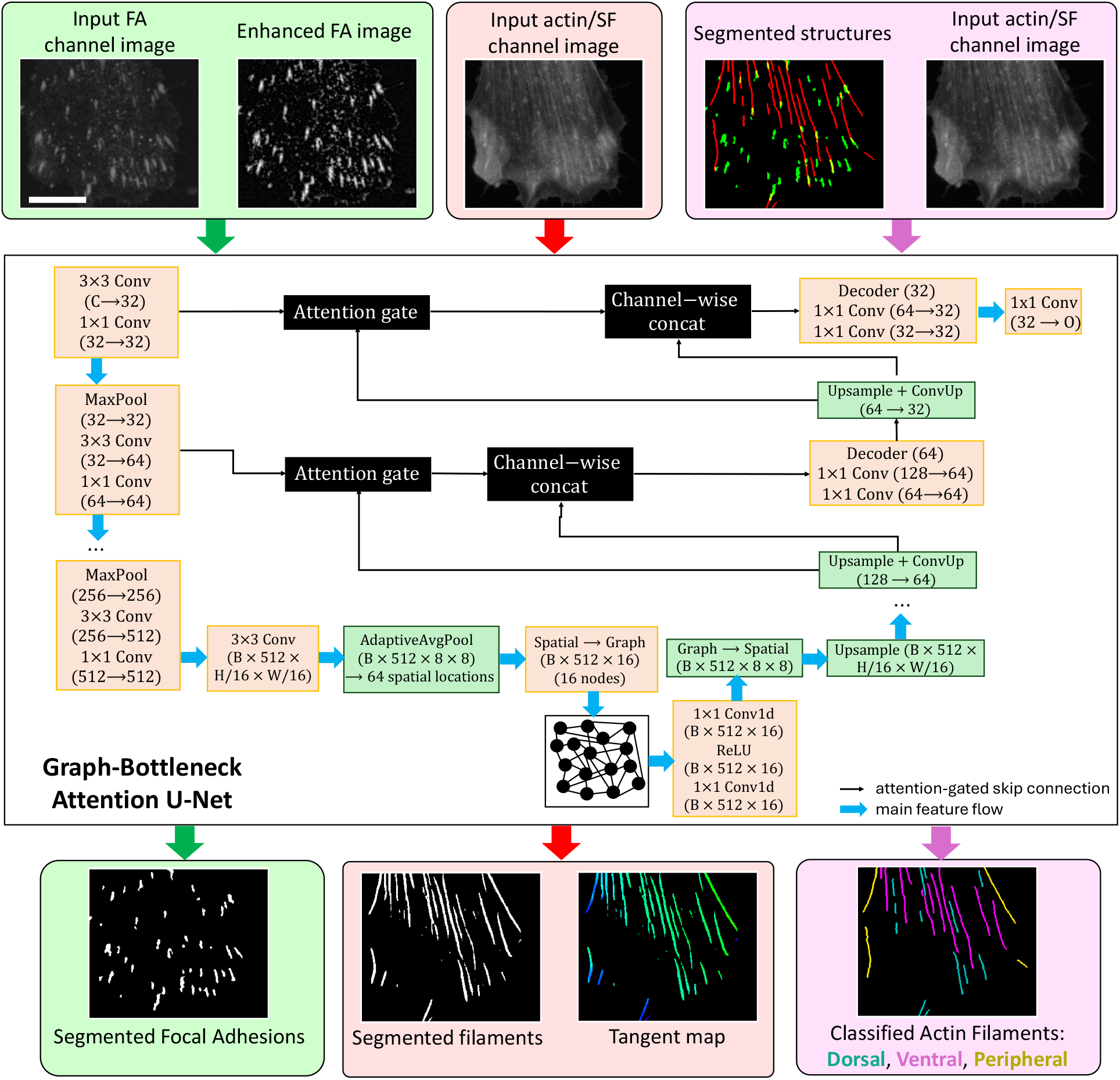
Task-specific Graph-Bottleneck Attention U-Nets support the three FilAdh branches. The FA model maps the raw and NAFNet-GAN-enhanced FA channel images to a focal-adhesion mask. The SF model maps the actin/SF channel image to both a filament mask and an undirected tangent map. The classifier maps the actin/SF channel image together with the FA and SF masks to background, dorsal, ventral, and peripheral probabilities. Each model uses an encoder, a global graph bottleneck based on learned soft spatial-to-node assignments, attention-gated skip connections, and a task-specific decoder.

### FilAdh achieves the highest focal-adhesion overlap on the pooled test set

FilAdh was compared with four FA benchmark methods on the same 57 held-out images using the prespecified benchmark protocol and a common evaluation pipeline (Figure 6; Table 4).

**Table 4:**
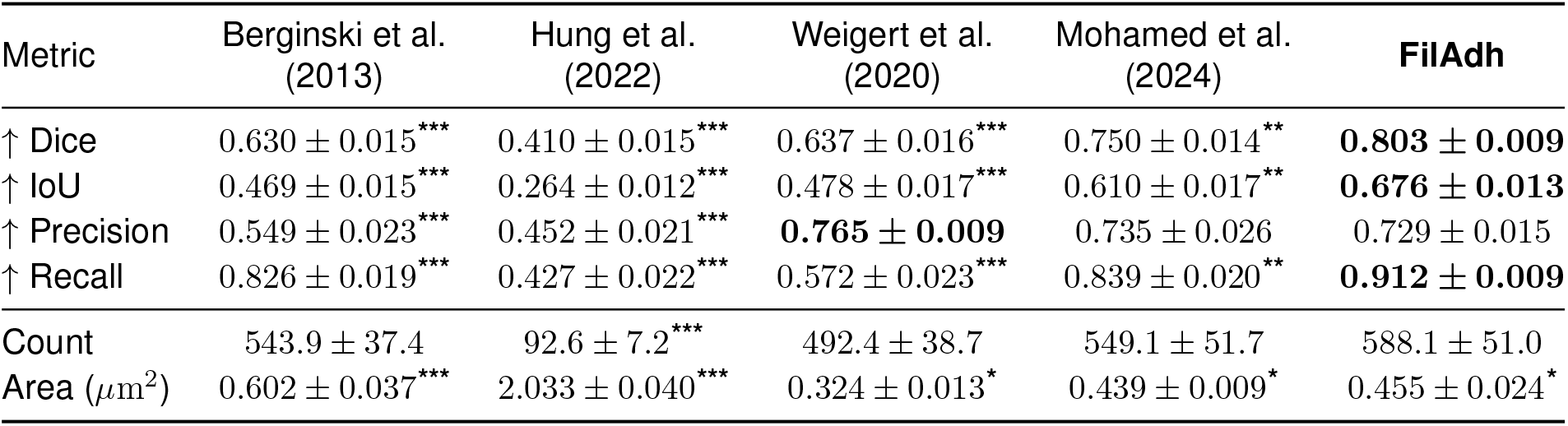
FA segmentation and morphometry on the clean held-out test set. Mean *±* s.e.m. across *n* = 57 held-out images. Bold marks the best observed segmentation mean. Asterisks indicate Holm-adjusted *p* values from two-sided paired *t* tests on image-level pairs: each compared method versus FilAdh for segmentation metrics and each prediction versus its corresponding reference for count and area. ^**\***^ *p <* 0.05, ^**\*\***^ *p <* 0.01, ^**\*\*\***^ *p <* 0.001.

| Metric | Berginski et al.<br>(2013) | Hung et al.<br>(2022) | Weigert et al.<br>(2020) | Mohamed et al.<br>(2024) | FilAdh |
| --- | --- | --- | --- | --- | --- |
| ↑ Dice | $0.630 \pm 0.015^{***}$ | $0.410 \pm 0.015^{***}$ | $0.637 \pm 0.016^{***}$ | $0.750 \pm 0.014^{**}$ | **$0.803 \pm 0.009$** |
| ↑ IoU | $0.469 \pm 0.015^{***}$ | $0.264 \pm 0.012^{***}$ | $0.478 \pm 0.017^{***}$ | $0.610 \pm 0.017^{**}$ | **$0.676 \pm 0.013$** |
| ↑ Precision | $0.549 \pm 0.023^{***}$ | $0.452 \pm 0.021^{***}$ | **$0.765 \pm 0.009$** | $0.735 \pm 0.026$ | $0.729 \pm 0.015$ |
| ↑ Recall | $0.826 \pm 0.019^{***}$ | $0.427 \pm 0.022^{***}$ | $0.572 \pm 0.023^{***}$ | $0.839 \pm 0.020^{**}$ | **$0.912 \pm 0.009$** |
| Count | $543.9 \pm 37.4$ | $92.6 \pm 7.2^{***}$ | $492.4 \pm 38.7$ | $549.1 \pm 51.7$ | $588.1 \pm 51.0$ |
| Area ($\mu\text{m}^2$) | $0.602 \pm 0.037^{***}$ | $2.033 \pm 0.040^{***}$ | $0.324 \pm 0.013^*$ | $0.439 \pm 0.009^*$ | $0.455 \pm 0.024^*$ |

**Figure 6:**
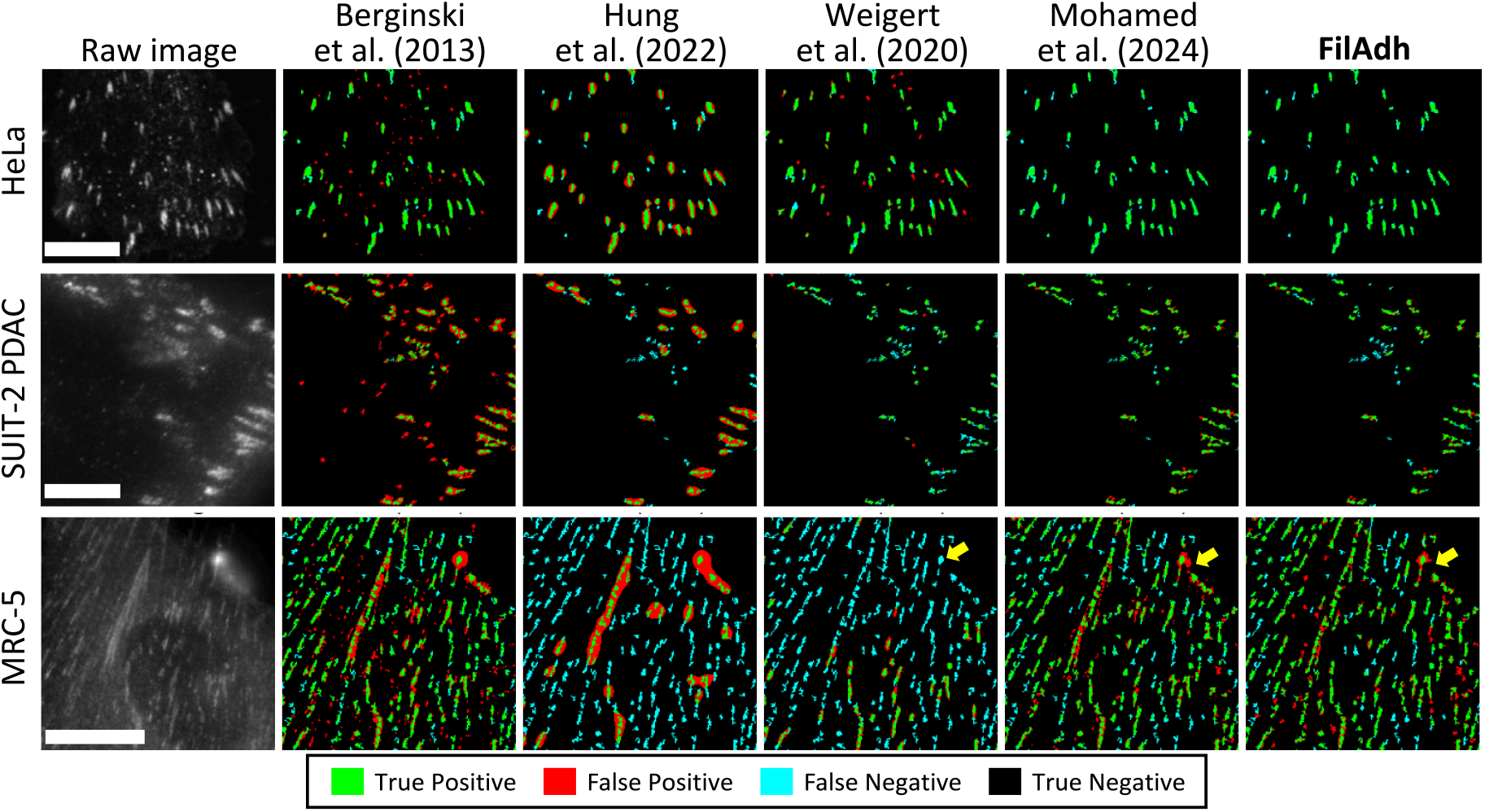
FA agreement on clean held-out images. Rows show HeLa, SUIT-2 PDAC, and MRC-5 examples; columns show raw FA channel image and benchmark predictions relative to the FA reference. Green denotes true positives, red false positives, cyan false negatives, and black true-negative background. Yellow arrows highlight the same challenging MRC-5 adhesion, showing undersegmentation by Weigert et al., oversegmentation by Mohamed et al., and closer reference agreement by FilAdh. Scale bars, 10 *µ*m.

Under this protocol, FilAdh achieved the highest Dice (0.803), IoU (0.676), and recall (0.912). In paired analyses of the 57 image-level values, all three metrics were significantly higher than those of every compared method after Holm adjustment (Table 4). Weigert et al. achieved the highest precision (0.765 versus 0.729 for FilAdh) but substantially lower recall (0.572), indicating a more conservative assignment of adhesion foreground.

The agreement maps in Figure 6 illustrate the practical effect of these differences: in the selected examples, FilAdh retained more weak adhesion regions while limiting boundary expansion. These examples are illustrative; the quantitative conclusions come from the image-level comparisons with the expert-curated references. Supplemental Figures S1–S3 show the results by dataset. The largest pooled differences were concentrated in the HeLa and SUIT-2 cohorts, while the smaller MRC-5 cohort showed metric-dependent rankings and wider uncertainty.

### FilAdh surpasses classical methods and performs comparably to deep learning methods on stress-fiber topology

FilAdh was compared with four SF benchmark methods on the fixed held-out images; the deep-learning comparators were retrained using the shared splits and binary SF references (Figure 7; Table 5).

**Table 5:**
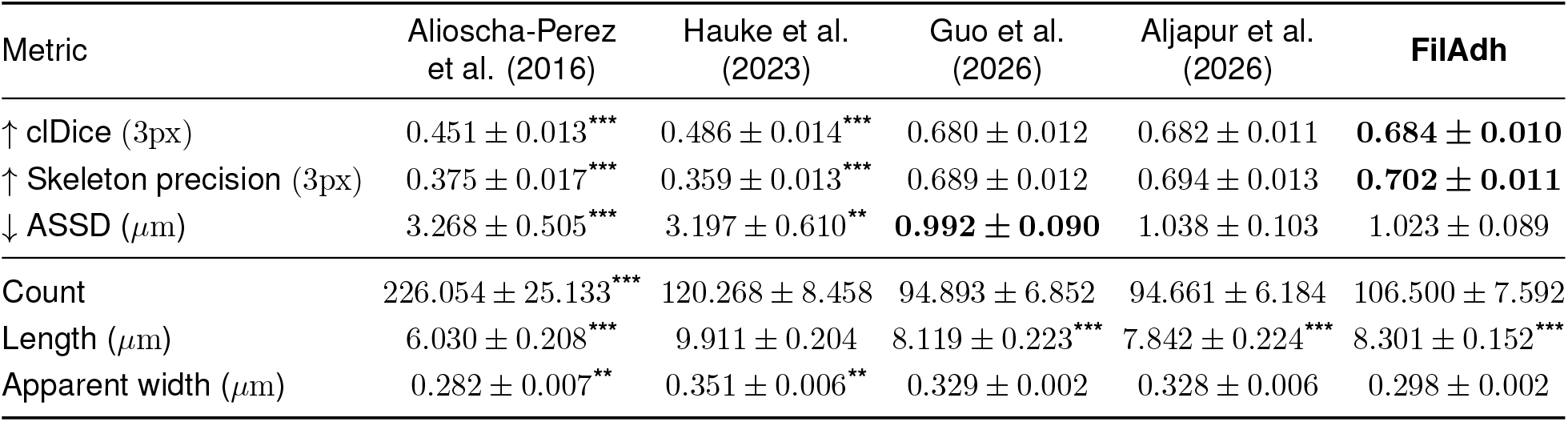
SF segmentation and morphometry on the clean held-out test set. Mean *±* s.e.m. across *n* = 57 held-out images. Bold marks the best observed topology mean. Asterisks indicate Holm-adjusted *p* values from two-sided paired *t* tests on image-level pairs: each compared method versus FilAdh for topology metrics and each prediction versus its corresponding reference for count, length, and apparent width. ^**\***^ *p <* 0.05, ^**\*\***^ *p <* 0.01, ^**\*\*\***^ *p <* 0.001.

**Figure 7:**
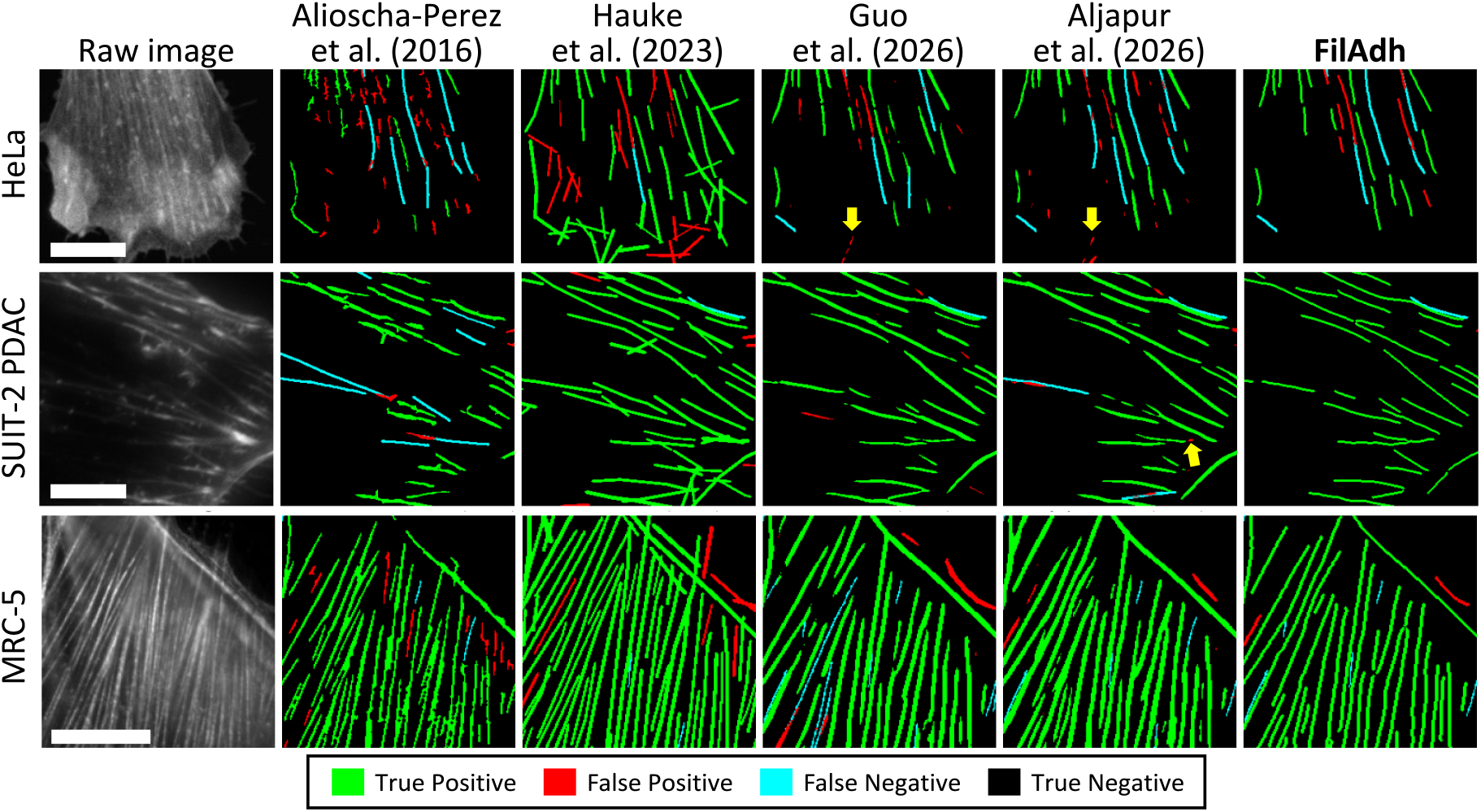
SF topology agreement on clean held-out images. Rows show HeLa, SUIT-2 PDAC, and MRC-5 examples; columns show raw actin/SF channel images and benchmark predictions relative to the expert-curated SF reference. Green denotes matched filament regions, red false-positive predictions, cyan false-negative reference regions, and black true-negative background. Yellow arrows highlight representative false-positive segments retained by the learned comparators but not present in the FilAdh reconstruction. Scale bars, 10 *µ*m.

FilAdh achieved the highest clDice (0.684) and skeleton precision (0.702). Both metrics were significantly higher than those of the two classical methods after Holm adjustment, whereas SynSeg and FAST produced comparable image-level results (Table 5). SynSeg had the lowest ASSD (0.992 versus 1.023 *µ*m for FilAdh), although this difference was not statistically significant after Holm adjustment. The three learned methods therefore performed within a narrow range under the present benchmark; this should not be interpreted as evidence of equivalence.

The examples in Figure 7 show local reconstruction errors that are not fully captured by the pooled topology metrics. Yellow arrows mark short false-positive segments retained by SynSeg or FAST but removed by FilAdh, while other regions show small centerline deviations from the expert-curated reference. These examples help explain the error patterns behind the pooled scores; the formal method comparisons remain those reported in Table 5.

Morphometric analysis revealed opposing consequences of reconstruction error. Fragmentation increased filament counts and shortened reconstructed objects, whereas merging or excessive reconnection reduced counts and increased apparent length. Dataset-stratified analyses showed that surface-distance and morphometric agreement varied with acquisition context (Supplemental Figures S4–S6).

### Degradation reveals distinct failure modes in adhesions and stress fibers

Degradation-aware checkpoints were evaluated on the same 57 held-out test images under four matched conditions: clean input, Poisson-Gaussian noise, Gaussian defocus, and point-spread-function (PSF) blur. These synthetic conditions serve as matched stress tests of specific acquisition-related failures and do not estimate how frequently such failures occur in routine microscopy. The clean columns in Tables 6 and 7 therefore report the performance of the degradation-aware checkpoints on clean input; they do not correspond to the clean-image-optimized checkpoints reported above.

**Table 6:**
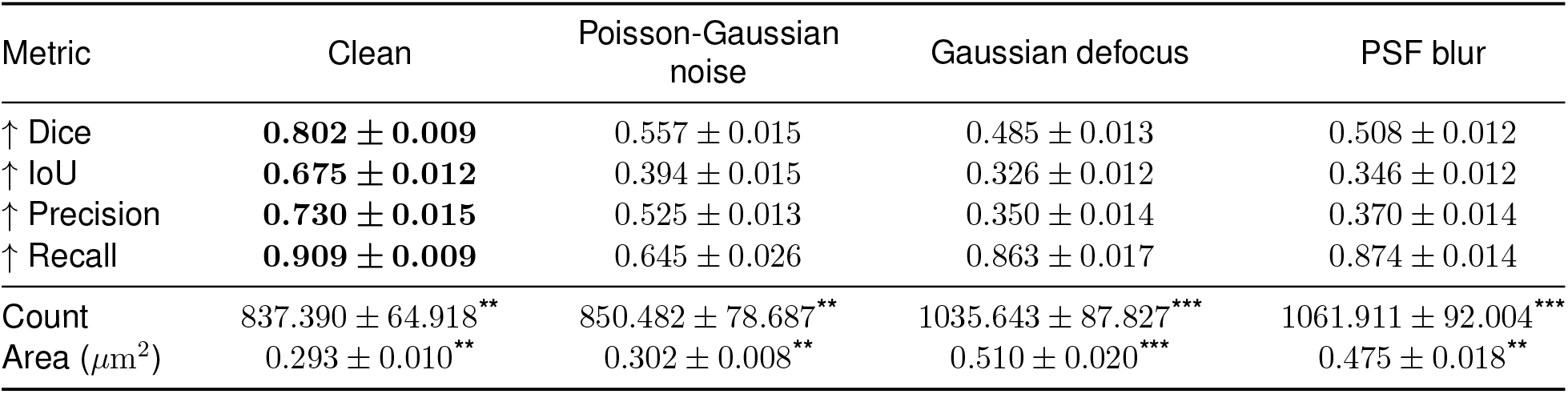
Degradation-aware FA performance across matched test conditions. Mean *±* s.e.m. across *n* = 57 matched images. Bold marks the best observed mean and is descriptive. Asterisks indicate Holm-adjusted *p* values from two-sided paired *t* tests comparing each image-level prediction with its corresponding reference for count and area. No inferential comparisons were performed between degradation conditions. ^**\***^ *p <* 0.05, ^**\*\***^ *p <* 0.01, ^**\*\*\***^ *p <* 0.001. Conditions are repeated within images.

| Metric | Clean | Poisson-Gaussian noise | Gaussian defocus | PSF blur |
| --- | --- | --- | --- | --- |
| ↑ Dice | **$0.802 \pm 0.009$** | $0.557 \pm 0.015$ | $0.485 \pm 0.013$ | $0.508 \pm 0.012$ |
| ↑ IoU | **$0.675 \pm 0.012$** | $0.394 \pm 0.015$ | $0.326 \pm 0.012$ | $0.346 \pm 0.012$ |
| ↑ Precision | **$0.730 \pm 0.015$** | $0.525 \pm 0.013$ | $0.350 \pm 0.014$ | $0.370 \pm 0.014$ |
| ↑ Recall | **$0.909 \pm 0.009$** | $0.645 \pm 0.026$ | $0.863 \pm 0.017$ | $0.874 \pm 0.014$ |
| Count | $837.390 \pm 64.918^{**}$ | $850.482 \pm 78.687^{**}$ | $1035.643 \pm 87.827^{***}$ | $1061.911 \pm 92.004^{***}$ |
| Area ($\mu\text{m}^2$) | $0.293 \pm 0.010^{**}$ | $0.302 \pm 0.008^{**}$ | $0.510 \pm 0.020^{***}$ | $0.475 \pm 0.018^{**}$ |

**Table 7:**
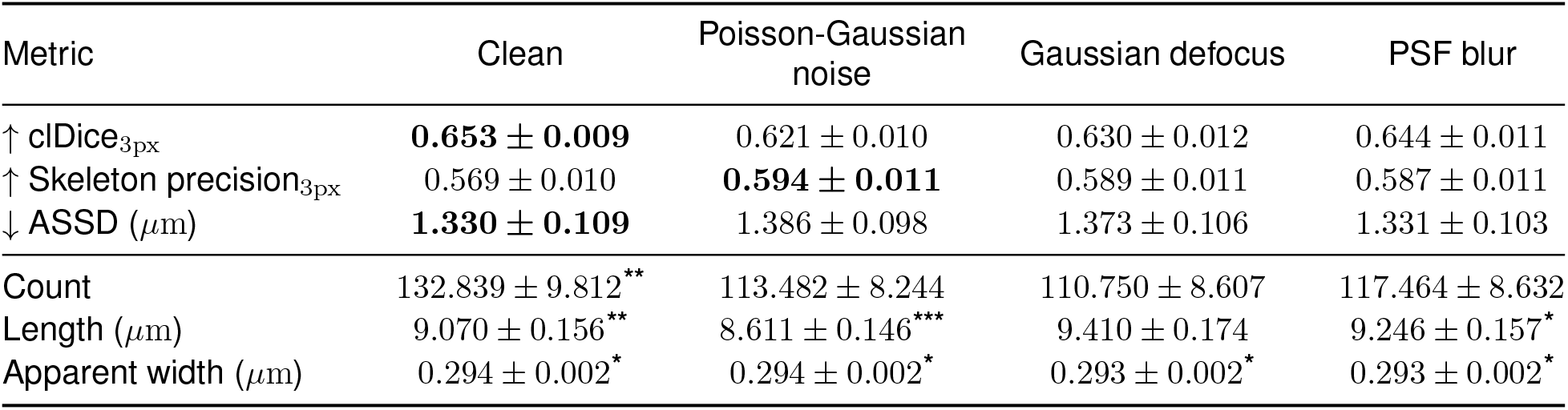
Degradation-aware SF performance across matched test conditions. Mean *±* s.e.m.; *n* = 57 test images. Bold marks the best observed mean and is descriptive. Asterisks indicate Holm-adjusted *p* values from two-sided paired *t* tests comparing each image-level prediction with its corresponding reference for count, length, and apparent width. No inferential comparisons were performed between degradation conditions. ^**\***^ *p <* 0.05, ^**\*\***^ *p <* 0.01, ^**\*\*\***^ *p <* 0.001. Conditions are repeated within images.

The observed FA errors differed across the tested degradations. Gaussian defocus and PSF blur broadened adhesion boundaries and merged closely spaced objects, increasing false-positive area and producing the largest observed reductions in precision, Dice, and IoU (Figure 8; Table 6). Poisson-Gaussian noise produced a different pattern: weak adhesion signal was lost, false-negative regions increased, and recall decreased while the boundaries of retained objects remained comparatively well defined. Degradation-aware training did not eliminate either error pattern completely, and blur continued to produce excess foreground.

**Figure 8:**
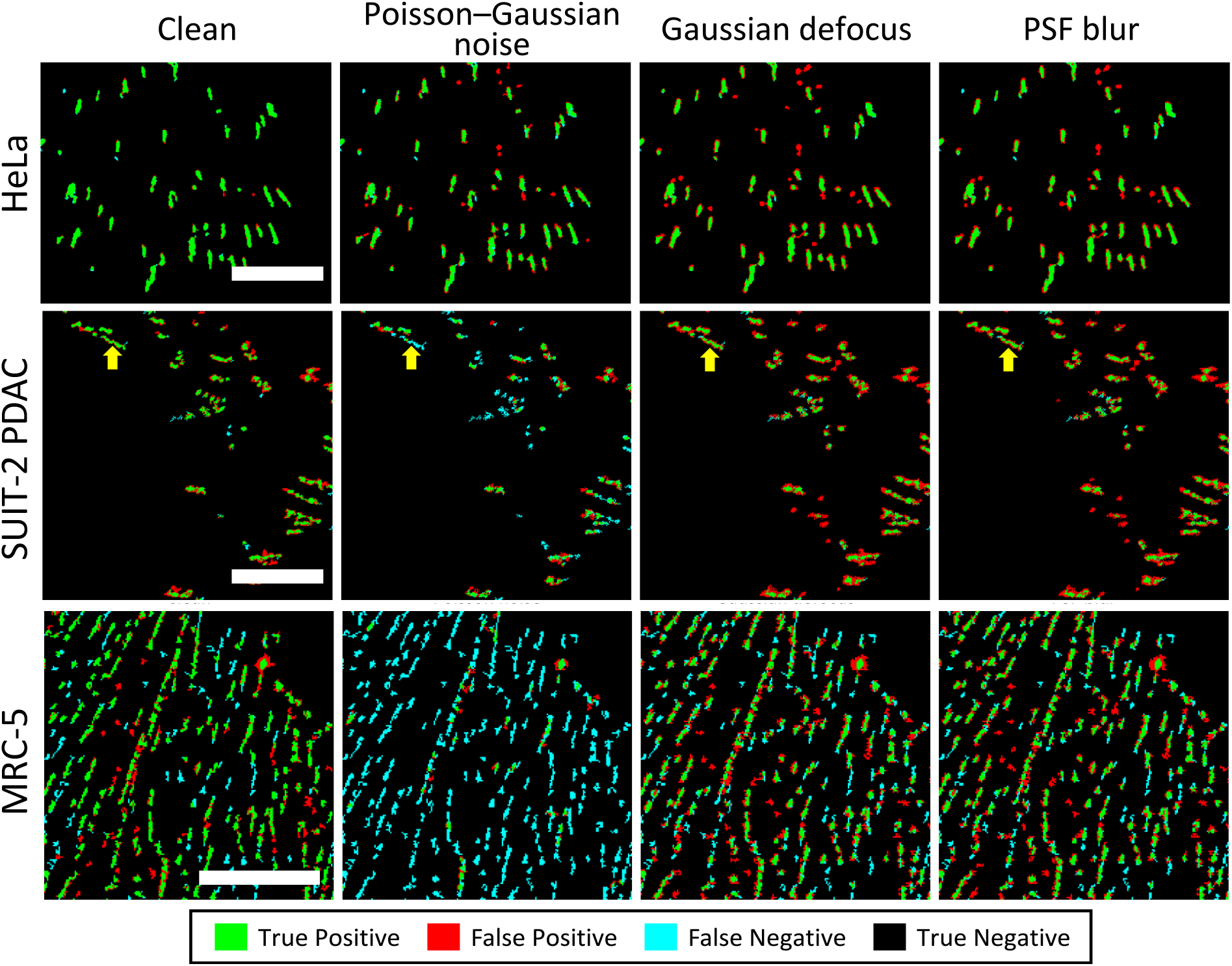
FA errors under matched synthetic degradations. Rows show HeLa, SUIT-2 PDAC, and MRC-5 examples; columns show clean input, Poisson-Gaussian noise, Gaussian defocus, and PSF blur. All predictions are compared with the unchanged FA reference. Green denotes true positives, red false positives, cyan false negatives, and black true-negative background. Yellow arrows track the same representative SUIT-2 adhesion across conditions and illustrate a change in error type: Poisson-Gaussian noise increases the cyan reference area missed by the prediction, whereas Gaussian defocus and PSF blur increase the red predicted area extending beyond the reference. Scale bars, 10 *µ*m.

Dataset-stratified analyses showed that the relative contribution of missed and enlarged adhesions depended on acquisition context (Supplemental Figures S7–S9). Recall loss under Poisson-Gaussian noise was most pronounced in lower-signal images, whereas both blur models shifted per-image area distributions toward larger adhesions, particularly in datasets containing densely packed neighboring structures.

SF topology was more stable across the tested conditions than FA overlap. clDice decreased from 0.653 on clean images to 0.621 under Poisson-Gaussian noise, while the observed changes in ASSD and skeleton precision were smaller (Figure 9; Table 7). The agreement maps and qualitative examples show that degradation mainly removed weak branches or introduced short local breaks rather than disrupting the filament network as a whole, although low-contrast branches remained vulnerable.

**Figure 9:**
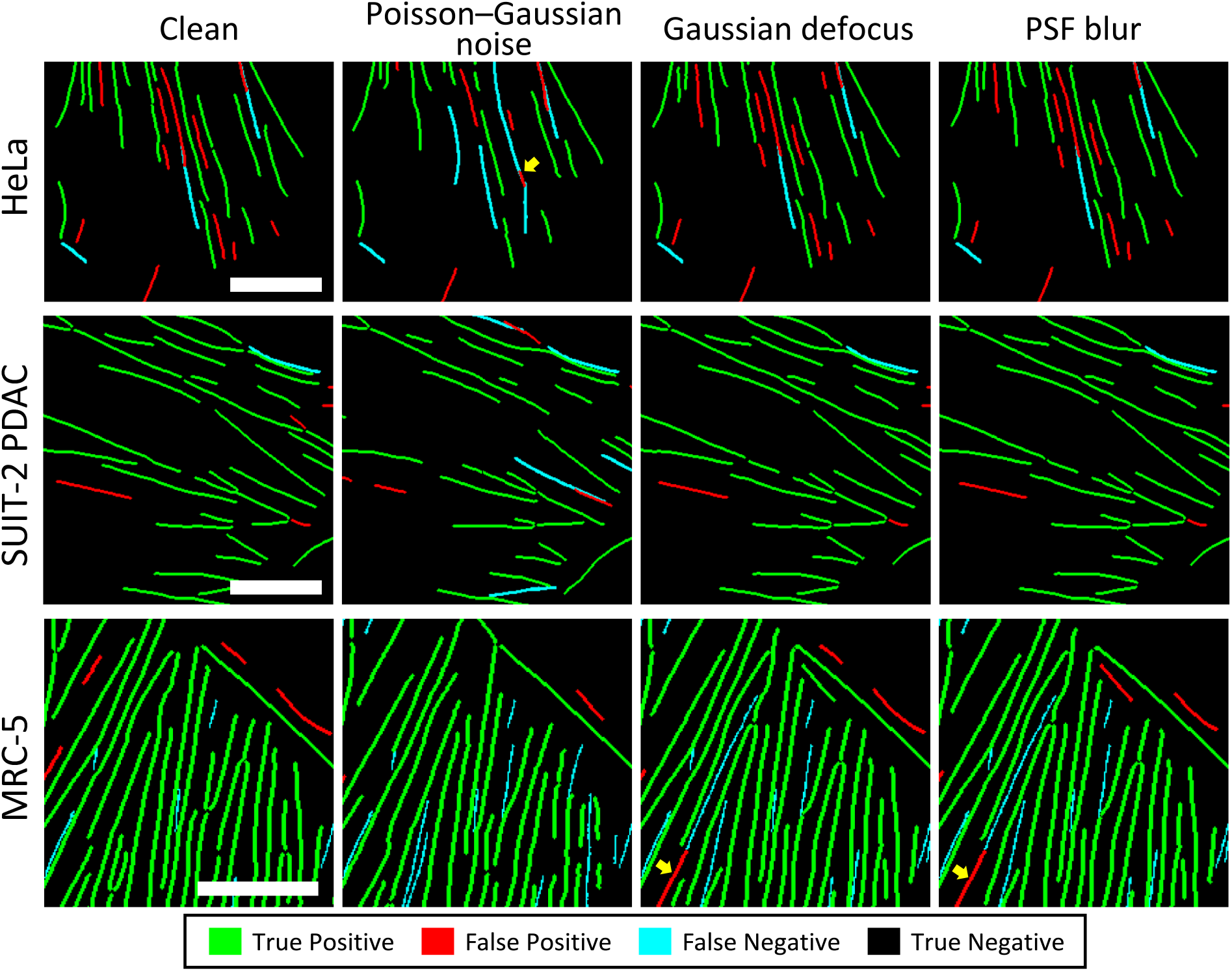
SF topology under matched synthetic degradations. Rows show HeLa, SUIT-2 PDAC, and MRC-5 examples; columns show clean input, Poisson-Gaussian noise, Gaussian defocus, and PSF blur. All predictions are compared with the unchanged SF reference. Green denotes true positives, red false positives, cyan false negatives, and black true-negative background. Yellow arrows highlight representative filament-level errors. The HeLa arrow under Poisson-Gaussian noise marks a reference centerline segment missed (cyan), whereas the MRC-5 arrows under Gaussian defocus and PSF blur mark additional predicted centerline segments absent from the reference (red). Scale bars, 10 *µ*m.

Dataset-stratified analyses showed that this relative topological stability did not eliminate morphometric bias (Supplemental Figures S10–S12). Length distributions remained comparatively close to the reference in SUIT-2, whereas larger count and apparent width offsets were observed in HeLa and MRC-5. These results underscore the need to interpret topology metrics together with object-level measurements.

### Conditional SF classification using expert-curated masks exceeds 80% accuracy

We evaluated classification using the actin/SF channel image together with the expert-curated FA and SF masks. This ensured that every prediction was assigned to a known filament and allowed us to measure class assignment independently of upstream reconstruction errors. Using predicted filaments at this stage would confound classification errors with reconstruction errors—fragmentation, merging, and false positives—because a predicted object may not correspond uniquely to a reference filament. We therefore accumulated the pixel-wise class probabilities within each expert-curated SF instance and assigned one label by object-level voting.

Object-level accuracy was 87.3% for HeLa, 83.0% for SUIT-2, and 81.5% for MRC-5 (Figure 10). Most errors involved dorsal–ventral confusion, whereas the MRC-5 results also showed dorsal–peripheral confusion. These accuracies were summarized descriptively because multiple filament instances originated from each source image and were not treated as independent biological replicates.

**Figure 10:**
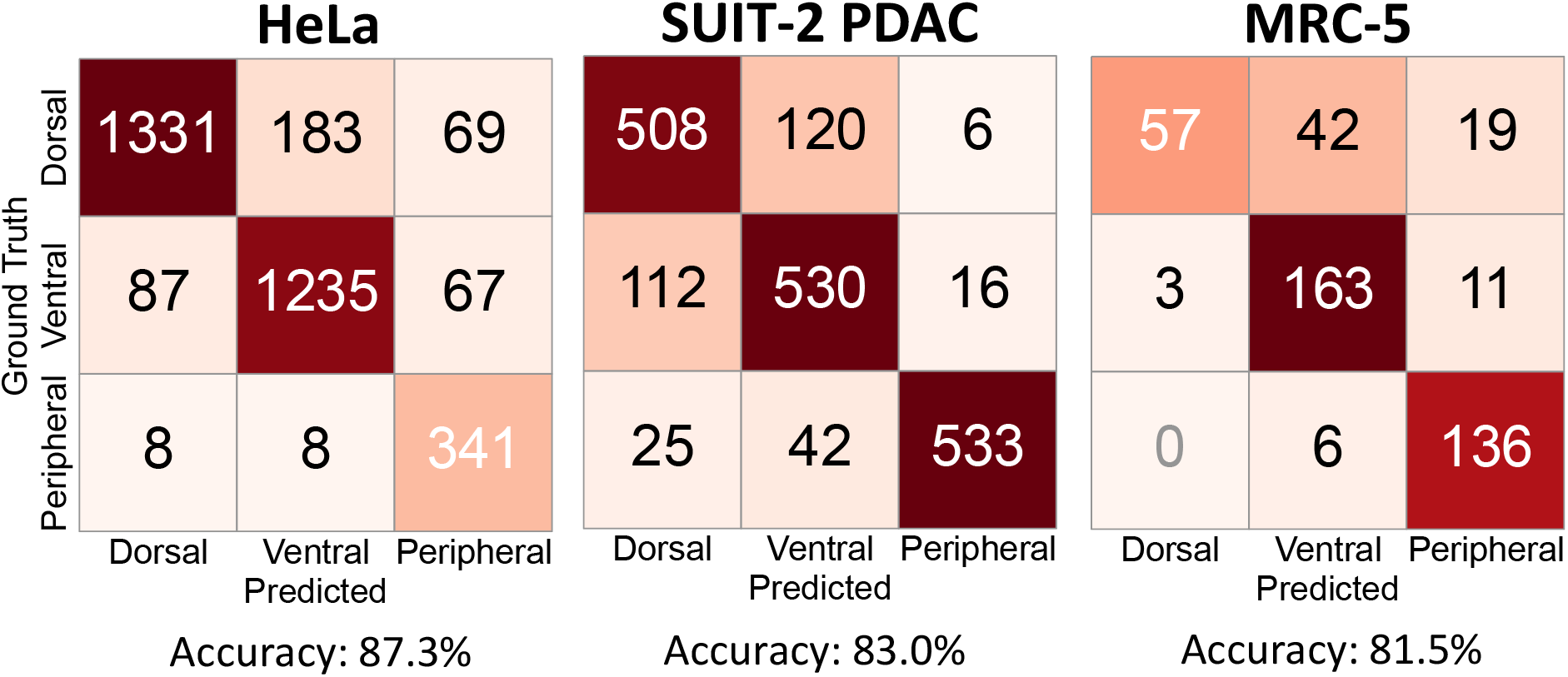
Conditional classification of expert-curated SF instances. Confusion matrices show object-level predictions obtained using the actin/SF channel image and expert-curated FA and SF masks. Values are raw counts of reference filament instances. Accuracy was 87.3% for HeLa (3,329 filaments from 30 images), 83.0% for SUIT-2 (1,892 filaments from 17 images), and 81.5% for MRC-5 (437 filaments from 10 images). Fila-ment instances were not treated as independent biological replicates.

### Hardware-matched segmentation-network inference times

Segmentation-network inference efficiency was characterized on the same fixed heldout subset of 57 images used for the biological evaluations. All locally timed networks were evaluated on an AMD Ryzen 7 5800X central processing unit (CPU) and an NVIDIA GeForce RTX 5060 Ti graphics processing unit (GPU) with 16 GB of memory. Timing started after model loading and one warm-up pass and covered the interval from image loading through forward inference. These measurements provide a hardware-matched comparison of the neural segmentation stages and do not represent complete pipeline wall-clock time.

Across all three native image sizes, the 2.44-million-parameter FilAdh Graph-Bottleneck Attention U-Net had the shortest observed GPU and CPU inference times among the locally benchmarked networks (Table 8). FA segmentation was compared with the 26.07-million-parameter U-Net++ ResNet34 implementation of Mohamed et al.^17,19^, whereas SF segmentation was compared with the 13.21-million-parameter SynSeg U-Net and the 26.07-million-parameter FAST implementation^24,25^. These network-only measurements exclude FA enhancement and watershed, SF instance reconstruction, output generation, and morphometric analysis.

**Table 8:** Native-resolution segmentation-network inference time. Values are mean *±* standard error of the mean (s.e.m.) seconds per image after one warm-up pass (*n* = 10 MRC-5, *n* = 30 HeLa, and *n* = 17 SUIT-2 images). Network-level timings exclude FA enhancement and watershed separation, SF instance reconstruction, output generation, and morphometric analysis. Bold denotes the shortest observed mean within each image set and task. Runtime measurements were descriptive, and no inferential comparisons were performed.

| Task | Image set | Network and source | Parameters | GPU (s) | CPU (s) |
| --- | --- | --- | --- | --- | --- |
| Focal-adhesion segmentation | MRC-5, $520 \times 696$ | **FilAdh Graph-Bottleneck Attention U-Net** | 2.44M | **0.017 ± 0.000** | **0.674 ± 0.011** |
| | | U-Net++ ResNet34, Mohamed et al.<sup>17,19</sup> | 26.07M | 0.032 ± 0.001 | 0.714 ± 0.013 |
| | HeLa, $1600 \times 1600$ | **FilAdh Graph-Bottleneck Attention U-Net** | 2.44M | **0.094 ± 0.000** | **4.657 ± 0.021** |
| | | U-Net++ ResNet34, Mohamed et al.<sup>17,19</sup> | 26.07M | 0.140 ± 0.001 | 4.964 ± 0.035 |
| | SUIT-2, $2560 \times 2160$ | **FilAdh Graph-Bottleneck Attention U-Net** | 2.44M | **0.214 ± 0.002** | **10.215 ± 0.049** |
| | | U-Net++ ResNet34, Mohamed et al.<sup>17,19</sup> | 26.07M | 0.320 ± 0.003 | 11.016 ± 0.008 |
| Stress-fiber segmentation | MRC-5, $520 \times 696$ | **FilAdh Graph-Bottleneck Attention U-Net** | 2.44M | **0.0190 ± 0.0003** | **0.696 ± 0.002** |
| | | SynSeg U-Net, Guo et al.<sup>24</sup> | 13.21M | 0.0312 ± 0.0004 | 1.675 ± 0.013 |
| | | U-Net++ ResNet34, Aljapur et al.<sup>25</sup> | 26.07M | 0.0275 ± 0.0003 | 0.767 ± 0.007 |
| | HeLa, $1600 \times 1600$ | **FilAdh Graph-Bottleneck Attention U-Net** | 2.44M | **0.1017 ± 0.0014** | **5.132 ± 0.006** |
| | | SynSeg U-Net, Guo et al.<sup>24</sup> | 13.21M | 0.2324 ± 0.0019 | 12.911 ± 0.083 |
| | | U-Net++ ResNet34, Aljapur et al.<sup>25</sup> | 26.07M | 0.1401 ± 0.0018 | 5.273 ± 0.058 |
| | SUIT-2, $2560 \times 2160$ | **FilAdh Graph-Bottleneck Attention U-Net** | 2.44M | **0.2095 ± 0.0025** | **10.560 ± 0.154** |
| | | SynSeg U-Net, Guo et al.<sup>24</sup> | 13.21M | 0.5025 ± 0.0071 | 29.340 ± 0.361 |
| | | U-Net++ ResNet34, Aljapur et al.<sup>25</sup> | 26.07M | 0.3062 ± 0.0032 | 10.903 ± 0.158 |

Weigert et al.^16^ were excluded from runtime comparison because their TensorFlow implementation was executed in a different hardware and software environment.

## DISCUSSION

Focal adhesions and stress fibers are mechanically coupled, but they are usually quantified separately. This study shows that compact adhesions and extended filament networks can be analyzed within one workflow without treating them as the same geometric problem. FilAdh achieved the highest FA Dice, IoU, and recall in the benchmark, while its SF topology results fell within the same range as the deep-learning comparators (Tables 4 and 5). Using expert-curated FA and SF masks, it also classified dorsal, ventral, and peripheral fibers with 81.5%–87.3% accuracy. The degradation experiments revealed a different picture of robustness: SF topology remained relatively stable across the tested conditions, whereas FA measurements were more sensitive to noise and blur.

In both structures, changes in segmentation or reconstruction did not translate directly into changes in morphometry (Tables 6 and 7).

### Object-level measurements reveal the consequences of segmentation error

Segmentation scores and biological measurements were not interchangeable. Under Gaussian defocus, FA Dice decreased from 0.802 to 0.485, while mean predicted adhesion area increased from 0.293 to 0.510 *µ*m^2^ and count increased from 837.4 to 1035.6 per image (Table 6). Thus, blur-related foreground expansion can affect both apparent adhesion size and number, not only pixel overlap.

The SF measurements behaved differently. Across the degradation conditions, topology changed comparatively little, yet object measurements still shifted: mean filament count changed from 132.8 on clean inputs to 110.8 under Gaussian defocus, while mean length changed from 9.07 to 9.41 *µ*m (Table 7). These descriptive changes illustrate why FA overlap and SF topology scores should be interpreted alongside count, area, length, and apparent width rather than reduced to a single performance ranking.

### Compact adhesions and filamentous networks fail differently

The degradation experiments revealed a geometry-specific contrast. For FAs, Poisson-Gaussian noise reduced recall to 0.645, consistent with loss of weak adhesions, whereas Gaussian defocus and PSF blur preserved recall at 0.863 and 0.874 but reduced precision to 0.350 and 0.370, respectively (Figure 8; Table 6). The latter pattern reflects the boundary broadening and spurious foreground visible in the agreement maps.

SF topology was less variable over the tested ranges. Across the three degradations, clDice ranged from 0.621 to 0.644 and ASSD from 1.331 to 1.386 *µ*m (Figure 9; Table 7). The qualitative examples show localized missing or additional centerline segments rather than global loss of the filament network. The extended geometry of SFs was therefore associated with greater metric stability under the tested degradations than the compact, closely spaced geometry of FAs, although weak branches remained vulnerable.

Poisson-Gaussian noise approximates low-photon conditions associated with weak signal, short exposure, or photobleaching, whereas defocus and PSF blur approximate focal-plane error and optical broadening^11,30^. These controlled tests characterize scenario-specific sensitivities and do not estimate the expected frequency of such artifacts in routine microscopy. Signal level, focus, and optical blur should therefore be checked before automated morphometry is interpreted.

### Benchmark performance is strong but bounded by reference design

The benchmark provides its strongest evidence for FA segmentation, while showing that FilAdh performs comparably to the deep learning methods for SF topology. Reference construction is important for interpreting these rankings. The FA references were initialized using an image-adjusted Horzum workflow and subsequently corrected by experts, while the NAFNet-GAN training target was derived from the pre-threshold grayscale output of the same workflow (Table 1; Figure 4)^14^. This shared preprocessing lineage may favor agreement with the curated masks. However, enhancement was trained on grayscale targets, whereas segmentation was evaluated on held-out binary references after image-specific adjustment and collaborative correction. The FA scores should therefore be interpreted as agreement with an expert-curated reference standard rather than validation against an absolute ground-truth representation of FA morphology.

Benchmark implementation also affects the rankings. The quantitative benchmarked Horzum method was the fixed-parameter containerized implementation of Hung et al., rather than the image-adjusted procedure used during reference construction^15^. The SF references were manually traced and remain subject to optical resolution and annotator judgment. SynSeg and FAST were retrained on the common splits and references, whereas the classical SF configurations were selected using validation images and then fixed before held-out evaluation (Tables 1 and 2). The reported rankings are therefore conditional on the reference annotations, evaluated implementations, and prespecified parameter-selection protocol.

### Anchorage labels are two-dimensional structural phenotypes

Dorsal, ventral, and peripheral labels were assigned from the observed number of FA contacts, the position of the filament within the cell, and its orientation relative to the nearest apparent cell boundary. These labels describe two-dimensional structural phenotypes in the analyzed image plane and do not directly establish complete three-dimensional connectivity or force transmission. This distinction is especially important for SUIT-2, which was acquired using TIRF microscopy. TIRF preferentially excites fluorophores near the coverslip, providing strong contrast for basal FAs while potentially omitting portions of an SF that curve away from the substrate^31–33^. FA endpoints may therefore remain visible while an intervening filament segment extends beyond the TIRF evanescent field, creating an apparent break and potentially altering the assigned two-dimensional class.

The reported labels should therefore be interpreted as two-dimensional, image-plane structural phenotypes rather than direct measurements of complete three-dimensional FA-SF connectivity. For TIRF data, these phenotypes are additionally biased toward structures located near the cell-substrate interface. Volumetric confocal imaging or another three-dimensional acquisition strategy would be required to distinguish true loss of connectivity from limited axial sampling.

### Limitations of the study

Fixed held-out partitions prevented test-set leakage, while pooled training and dataset-balanced sampling reduced dominance of the larger HeLa collection during model fitting. Nevertheless, repetition within the training and validation loaders does not increase biological diversity, and the relatively small SUIT-2 and MRC-5 cohorts limit the precision of dataset-specific performance estimates. Validation also remains restricted to the datasets analyzed in the present study. Independent cross-laboratory datasets will therefore be required to establish generalization across acquisition systems, staining protocols, and biological contexts.

The controlled Poisson-Gaussian noise, Gaussian defocus, and PSF blur experiments isolate specific acquisition-related failure modes, but they do not reproduce the full complexity of naturally degraded microscopy data, in which signal loss (low exposure and photobleaching), optical errors (defocus and aberrations), and channel misalignment may co-occur. Because the conditions were imposed synthetically rather than sampled from a prospective audit of microscope output, the present study cannot estimate their prevalence in routine experiments or define a universal image-quality acceptance threshold. The results instead identify error patterns that can arise when low signal, focus error, or optical broadening remains after acquisition quality control.

We also did not propagate these perturbations through the complete segmentation-to-classification pipeline, so their effect on final anchorage assignments remains unknown.

### Conclusions

Overall, FilAdh offers a practical way to study focal adhesions and stress fibers together while preserving the different information needed for each structure. A single model for each task was trained across all three cell lines and microscopy settings, with strong FA segmentation and SF topology comparable to the deep-learning benchmarks on held-out images. Using expert-curated masks, SF classification remained above 80% across all three datasets. The degradation experiments showed that robustness depended on what was being measured: SF topology remained relatively stable, whereas FA measurements were more sensitive to noise and blur. This distinction matters because good segmentation does not necessarily mean that the measurements derived from it are equally reliable.

## STAR METHODS

### RESOURCE AVAILABILITY

#### Lead contact

Further information and requests for resources should be directed to the lead contacts, Dariusz Lachowski and Arrate Muñoz-Barrutia.

#### Materials availability

This study did not generate new unique reagents.

#### Data and code availability

Code, trained weights, fixed split manifests, model configurations, and analysis scripts are available to editors and reviewers through a private anonymized repository; access instructions accompany the submission. Before publication, the reviewed release will be deposited in a permanent public repository with a persistent identifier and an explicit software license.

The original source microscopy images are subject to the conditions imposed by their providers and are therefore not redistributed in the review repository. Derived annotations, predictions, and per-image measurements are provided where permitted. The public record will identify the source and access conditions for every dataset that cannot be redistributed.

### METHOD DETAILS

#### Study design and datasets

The analysis used paired fluorescence images of HeLa cervical carcinoma cells, SUIT-2 pancreatic ductal adenocarcinoma cells, and MRC-5 human lung fibroblasts. Across datasets, we refer to the channel reporting focal-adhesion signal as the FA channel and the channel reporting actin stress fibers as the actin/SF channel. HeLa images were acquired by spinning-disk confocal microscopy,^26^ SUIT-2 images by total internal reflection fluorescence (TIRF) microscopy,^27^ and MRC-5 images by laser-scanning confocal microscopy^23^. TIRF preferentially excites fluorophores within a shallow region adjacent to the coverslip; it therefore emphasizes basal FAs but may not capture SF segments that extend farther from the substrate^31,32^.

Training, validation, and test lists were fixed before final evaluation and were shared by all trainable methods. All reported segmentation, morphometry, degradation, classification, and local runtime results were computed exclusively from the fixed held-out test subsets comprising 30 HeLa, 17 SUIT-2 PDAC, and 10 MRC-5 images. One pooled model was trained for each task and checkpoint family across the three datasets; no dataset-specific fine-tuning was used for the reported test results.

To reduce dominance of the larger HeLa collection while preserving split membership, SUIT-2 and MRC-5 entries were repeated within the training and validation loaders to effective sizes of 258 and 266, respectively, while HeLa contributed 267 entries. Repetition changed sampling frequency but did not create additional biological replicates. Augmentation was applied only to training data, and no held-out test image was used for model fitting, threshold selection, hyperparameter selection, or checkpoint selection.

Dataset-specific calibration was 0.108 *µ*m*/pixel* for HeLa and SUIT-2, and 0.122 *µ*m*/pixel* for MRC-5. Images were stored as 16-bit tagged image file format (TIFF) files. The exported files did not retain complete acquisition metadata for objective numerical aperture, detector model, or exposure time; these values were not inferred from the processed images.

#### Reference annotation

FA candidate masks were generated in Fiji using a workflow adapted from Horzum et al.^14^. Following the general processing strategy described therein, images underwent background subtraction, contrast-limited adaptive histogram equalization (CLAHE), Laplacian-of-Gaussian (Mexican-hat) filtering, normalization, nonlinear intensity transformation, thresholding, and labeling. Filter and threshold parameters were adjusted image by image until visible paxillin-positive structures were adequately captured, as judged by the annotators, with particular attention to separating neighboring adhesions.

SFs were traced manually in Labkit^28^. Annotators followed the visible centerline and width of each fiber through dense regions and retained a single instance across a weak gap when the local signal and geometry supported continuity. Ambiguous crossings, truncated fibers, and low-signal regions were reviewed using the paired actin and adhesion channels. Each traced SF was assigned a structural subtype according to its FA contacts, position within the cell, and orientation relative to the nearest apparent membrane. Dorsal SFs contacted an FA at one end and were oriented approximately perpendicular to the nearest apparent membrane. Ventral SFs contacted FAs at both ends, lay in the central region of the cell, and were oriented approximately tangent to the nearest apparent membrane. Peripheral SFs lay at the cell periphery and contacted FAs at both ends.

Candidate masks were reviewed collaboratively by G.R.-P. and M.H.-M., with annotation assistance from Beatriz Rabadán-Gallego. Ambiguous regions were discussed and corrected by consensus. This operator-guided procedure is reported explicitly because it introduces dependence on both preprocessing and expert judgment.

#### Benchmark implementation

All methods received the same held-out images at native resolution or at the resolution required by the released implementation, and predictions were returned to the reference grid before metric calculation. For FA analysis, an image-adjusted Horzum workflow contributed to reference construction, whereas quantitative benchmarking used the containerized Horzum implementation of Hung et al.^15^ with fixed parameters on the held-out test images.

For the classical SF methods, predefined grids covering the relevant detection, linking, and filtering parameters were evaluated on validation images. One configuration per method was selected from validation performance and frozen before held-out test evaluation. No test image or test reference was used for parameter selection. FilamentSensor 2.0, as reported by Hauke et al.^22^, was used as the current classical filament-analysis method; FAFCK^23^ is discussed as prior work combining FilamentSensor-based analysis with FA detection and cross-correlation. Classical outputs were converted to the common binary and instance-level evaluation representations.

SynSeg^24^ and FAST^25^ were not evaluated using pretrained outputs tied to their original datasets. Their released training frameworks were adapted to the present input format and retrained with the same training and validation splits and manual SF targets used for FilAdh. Published architecture and optimization settings were retained where compatible; changes were limited to input/output handling, target replacement, and conversion to the common evaluation representation. Validation data alone were used for checkpoint selection.

The Weigert et al. implementation was evaluated for segmentation accuracy in a compatible Google Colab environment because its TensorFlow stack is not compatible with the RTX 5060 Ti workstation used for local timing^16^. Runtime was not compared across these environments because both hardware and software differed. This exclusion applies only to runtime; the segmentation results remain part of the common held-out-image comparison.

#### NAFNet-GAN enhancement

The nonlinear-activation-free network–generative adversarial network (NAFNet-GAN) architecture and generator objective followed our previously described bioimage-restoration framework^20^. For the present FA application, the same architecture and loss formulation were retrained using raw FA channel images and Horzum-derived enhanced targets. The generator backbone followed NAFNet^34^. It used one input channel, one output channel, a base width of 16, encoder block depths of [2, 2, 4, 8], 12 middle NAFBlocks, and decoder block depths of [2, 2, 2, 2]. The PatchGAN-like discriminator used successive 4 *×* 4 convolutional blocks and produced a final local logit map^35^. Raw and target-enhanced images were min–max normalized, randomly cropped to 256 *×* 256 pixels, and augmented using synchronized horizontal and vertical flips.

Consistent with the previously reported NAFNet-GAN formulation, the generator objective combined learned perceptual image patch similarity (LPIPS), Charbonnier, Laplacian, frequency-domain, and adversarial terms^20^:

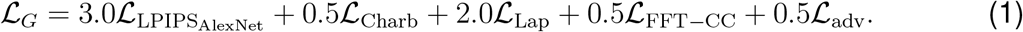

LPIPS compares images in a pretrained AlexNet feature space, whereas the Charbonnier term provides a smooth, robust pixel-wise approximation to absolute error^36,37^:

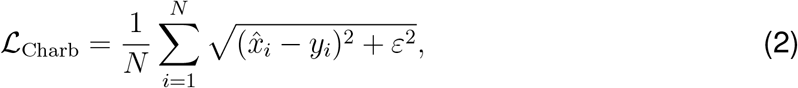

where 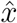 and *y* are the generated and target-enhanced images. The Laplacian term penalized disagreement between edge responses, and the fast Fourier transform (FFT) magnitude consistency loss term penalized disagreement in radially weighted frequency magnitudes. The adversarial generator term was 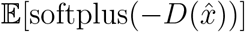; the discriminator used the hinge objective

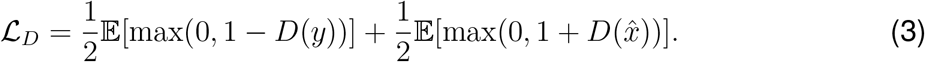

The discriminator was used only during training. R1 regularization was implemented but disabled in the reported configuration. Training used AdamW for 200 epochs with initial learning rate 5 *×* 10^−4^, cosine annealing with *T*_max_ = 40, batch size 8, random seed 42, and validation-loss checkpoint selection.

#### Graph-Bottleneck Attention U-Net

The encoder comprised four depthwise-separable convolution stages with 32, 64, 128, and 256 channels, with max pooling between stages. Each block used a depthwise 3 *×* 3 convolution, a pointwise 1 *×* 1 convolution, batch normalization, and a rectified linear unit. A deformable convolution mapped 256 channels to a 512-channel bottleneck^38,39^.

For global feature aggregation, adaptive average pooling reduced the bottleneck to an 8 *×* 8 grid. A learned 1 *×* 1 projection generated 16 soft assignment maps over the 64 pooled spatial locations, which aggregated the pooled features into 16 latent graph nodes. The features of each node were transformed by two 1 *×* 1 one-dimensional convolutions with an intervening rectified linear unit. The transformed node features were then redistributed to the spatial representation through the learned assignment maps, reshaped to the 8 *×* 8 grid, upsampled to the original bottleneck resolution, and fused residually with the convolutional features^40^. The decoder used bilinear upsampling, 1 *×* 1 channel reduction, attention-gated skip concatenation, and separable convolution blocks^41^.

The FA model received the raw and NAFNet-GAN-enhanced FA channel images as two input channels and produced a binary focal-adhesion logit map. The SF model received the actin/SF channel image and produced a binary filament logit map together with a two-channel undirected tangent representation, (cos 2*θ*, sin 2*θ*). This mask-and-tangent architecture was used for both the clean-image and degradation-aware SF checkpoints. The classification model received the actin/SF channel image, binary FA mask, and binary SF mask as three input channels and produced background, dorsal, ventral, and peripheral logits.

#### Training objectives and optimization

Task-specific objectives were used because the three branches present different optimization challenges: sparse FA foreground, thin and orientation-dependent SF topology, and imbalanced anchorage classes.

The FA segmentation objective combined Tversky and focal losses. Tversky loss was used to control the relative penalty assigned to missed adhesions and false-positive foreground in the presence of strong class imbalance. With soft true-positive, false-negative, and false-positive counts denoted by *TP, FN*, and *FP*,

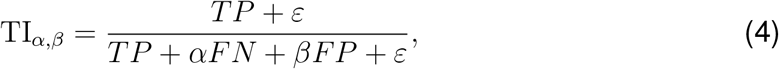

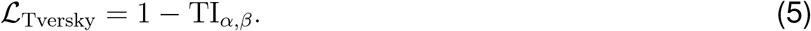

Focal loss was added to reduce the contribution of easily classified background pixels and emphasize harder foreground and boundary examples. The focal term was

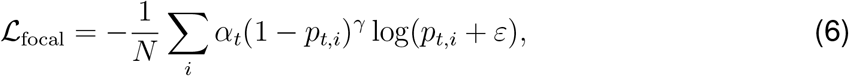

where *p*_*t,i*_ = *p*_*i*_ for foreground pixels and *p*_*t,i*_ = 1 *− p*_*i*_ for background pixels^42,43^. The combined FA objective was

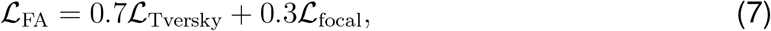

with Tversky parameters *α* = 0.9 and *β* = 0.1, focal exponent *γ* = 2, and *α*_*t*_ = 0.75 for foreground pixels and 0.25 for background pixels. The larger Tversky weight placed greater emphasis on recovering the sparse adhesion foreground, while the focal component retained sensitivity to difficult local errors.

Both SF checkpoints were trained to predict the binary filament mask and the undirected tangent representation. A dense tangent target was computed from the structure tensor of the binary mask after Gaussian smoothing with *σ* = 1 pixel^29^. Because filament orientation is undirected, the tangent orientation was encoded modulo *π* as

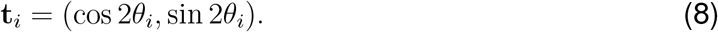

For foreground pixels *V* with valid local orientation,

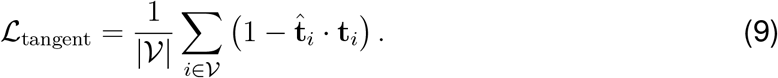

This cosine-based term penalized disagreement in local undirected orientation while treating directions separated by *π* as equivalent.

For both the clean-image and degradation-aware SF checkpoints, the binary filament mask used the same Tversky–focal formulation defined above:

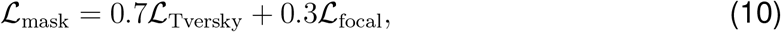

with Tversky parameters *α* = 0.9 and *β* = 0.1, focal exponent *γ* = 2, and *α*_*t*_ = 0.75 for foreground pixels and 0.25 for background pixels. The Tversky–focal mask loss emphasized recovery of the sparse filament foreground while retaining sensitivity to difficult local errors.

The complete SF objective, used unchanged for both checkpoint families, was

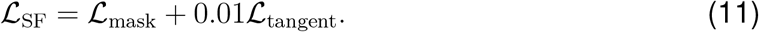

The tangent term provided auxiliary supervision for local filament orientation without dominating the primary mask-segmentation objective.

The filament classifier used weighted multiclass cross-entropy over background, dorsal, ventral, and peripheral labels:

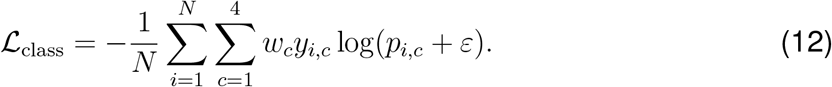

Class weighting reduced the influence of unequal class frequencies and prevented the most abundant structural labels from dominating optimization. The weights *w*_*c*_ were fixed from the training-set frequencies.

The clean-image FA segmentation checkpoint and the clean-image SF segmentation checkpoint used the same optimization settings. Each model was trained for 150 epochs with batch size 2 and 512 *×* 512 crops using AdamW with an initial learning rate of 5 *×* 10^−5^, weight decay 10^−6^, and cosine annealing with *T*_max_ = 30 and a minimum learning rate of 10^−6^.

The filament classifier was trained for 250 epochs with batch size 1 and 1024 *×* 1024 crops. Optimization used AdamW with an initial learning rate of 5 *×* 10^−4^ and weight decay 10^−5^.

The degradation-aware FA and SF checkpoints were initialized from their corresponding clean-image checkpoints and trained for 100 epochs with batch size 4 and 512 *×* 512 crops. Validation was performed on uncropped images. Both models were optimized using AdamW with an initial learning rate of 6 *×* 10^−4^, weight decay 10^−4^, and cosine annealing with warm restarts over four equal cycles and a minimum learning rate of 10^−6^.

For all models, the checkpoint with the lowest validation loss was retained for evaluation.

#### Inference and post-processing

The FA probability map was thresholded at 0.5. Adjacent foreground regions were separated by marker-controlled watershed on the negative Euclidean distance transform^44^.

Markers were local maxima with a minimum separation of 4 pixels; border maxima were permitted. If no marker was found, the binary component was retained after size filtering. Watershed regions smaller than 8 pixels were discarded, corresponding to approximately 0.09 µm^2^ in HeLa, 0.10 µm^2^ in SUIT-2, and 0.12 µm^2^ in MRC-5. Border-touching objects were retained when they passed the size criterion.

The SF probability map was thresholded before skeleton-based instance refinement. Tangent angles were treated as undirected orientations in [0, *π*). Paths with incompatible local orientation were separated, while opposing arms were reconnected only when both geometric direction and predicted tangent supported continuity. Labels were filtered by component size and expanded back onto the predicted foreground by constrained adaptive dilation. All thresholds were frozen before held-out evaluation and applied unchanged to the three datasets; exact values are reported in Supplemental Table S1.

For the reported conditional classification experiment, the classifier received the expert-curated binary SF foreground mask together with the expert-curated FA mask. The corresponding four-class SF annotation provided the pixel-wise target labels: background, dorsal, ventral, and peripheral. This configuration isolates anchorage-class assignment from errors in upstream FA and SF segmentation.

At inference, the network logits were converted to pixel-wise softmax probabilities. To obtain one prediction per reference filament, 8-connected components were extracted separately from the dorsal, ventral, and peripheral reference masks. No component dilation was applied. For each reference component *i* and non-background class *c ∈ C* = {dorsal, ventral, peripheral}, the class score was calculated by summing the corresponding probabilities over all pixels belonging to that component:

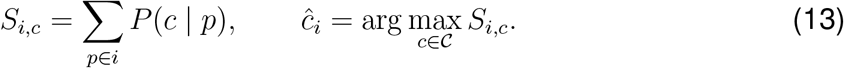

The class with the largest accumulated probability was assigned to the complete reference component. Background was excluded from the object-level vote because every evaluated component was defined by reference SF foreground. Because filament identity and spatial support were supplied by the reference annotation, no predicted-to-reference instance-matching step was required. Object-level accuracy was calculated as the number of correctly classified reference components divided by the total number of evaluated reference components.

#### Synthetic degradation

Images were converted to floating-point values, normalized as required, and clipped to [0, 1]. Degradation-aware training sampled clean images, point-spread-function (PSF) blur, Gaussian defocus, and Poisson–Gaussian noise, with the noise condition sampled more frequently to represent low-signal failure. Poisson–Gaussian noise was simulated by applying signal-dependent Poisson shot noise followed by additive, zero-mean Gaussian read noise:

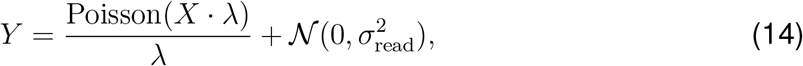

where *X* is the normalized clean image, *λ* is a peak-count scaling parameter, and *σ*_read_ is the read-noise standard deviation; the resulting image was clipped to [0, 1]. Both *λ* and *σ*_read_ were chosen artificially rather than calibrated to a specific camera. Validation used a deterministic version of the same schedule. At test time, each of the 57 held-out source images was evaluated once under each of the four matched conditions, yielding 57 image-level observations per condition. Reference masks remained unchanged across conditions, and repeated observations from the same source image were not treated as independent biological replicates. Parameter ranges, severity scaling, boundary handling, and sampling proportions are reported in Supplemental Table S2.

#### Runtime benchmarking

Runtime experiments were performed on the fixed held-out test subset comprising 30 HeLa, 17 SUIT-2 PDAC, and 10 MRC-5 images. Measurements were obtained on a workstation with an AMD Ryzen 7 5800X CPU and an NVIDIA GeForce RTX 5060 Ti GPU with 16 GB video memory. Model loading was completed before timing, one warmup inference preceded measurement, and GPU calls were synchronized around the timed region.

The timed region included image loading, normalization, padding, device transfer, forward inference, activation, and cropping. Output saving, visualization, instance postprocessing, and morphometric analysis were excluded. Runtime was measured independently for each source image at its native resolution and summarized within each dataset as mean *±* s.e.m.

For FA segmentation, 2.44-million-parameter Graph-Bottleneck Attention U-Net was compared with the 26.07-million-parameter U-Net++ ResNet34 baseline on the same held-out images. NAFNet-GAN enhancement and watershed separation were excluded from both the reported FilAdh network timing and the scope of the network-level comparison. For SF segmentation, the 2.44-million-parameter Graph-Bottleneck Attention UNet was compared with the 13.21-million-parameter SynSeg U-Net and the 26.07-million-parameter U-Net++ ResNet34 implementation used for FAST on the same held-out images. Orientation-guided instance refinement was excluded. Star-convex runtime was omitted because the hardware and software environments differed, as described above.

### QUANTIFICATION AND STATISTICAL ANALYSIS

#### Evaluation metrics and morphometry

FA masks were evaluated with Dice, IoU, precision, and recall. For predicted mask *P* and reference mask *R*,

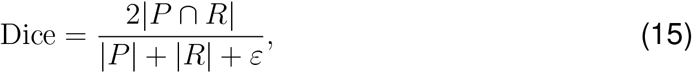

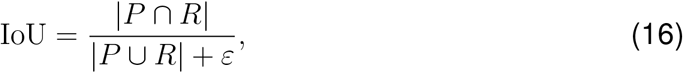

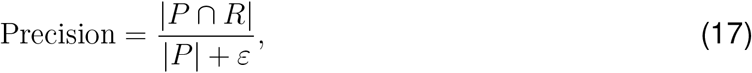

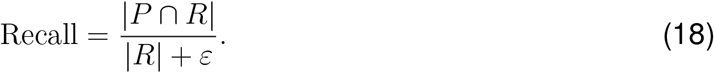

SF masks were skeletonized and evaluated with a 3-pixel tolerance. Skeleton precision was the fraction of predicted skeleton pixels within 3 pixels of the reference skeleton; skeleton sensitivity was defined analogously from reference to prediction. The tolerancebased clDice was their harmonic mean^45^.

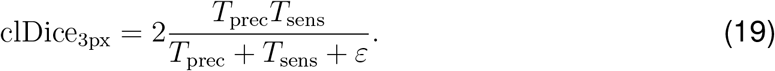

ASSD was calculated between the boundaries of the predicted and reference binary SF masks and converted to micrometers using dataset-specific calibration.

FA count was the number of retained watershed instances, and area was foreground pixel count multiplied by physical pixel area. For SFs, centerline length was the 8-connected geodesic skeleton length, and apparent width was component area divided by skeleton length. Counts and mean component measurements were calculated per image and then summarized across images. Empty masks contributed a count of zero; mean length and apparent width were omitted when no component remained.

#### Statistical analysis

Values are reported as mean *±* standard error of the mean (s.e.m.) unless otherwise stated. The source image, not an individual adhesion or filament, was the experimental unit. Duplicate rows sharing an image identifier were collapsed to one observation per method and source image.

All inferential comparisons used the same two-sided paired *t* test on matched image-level values. The paired contrast depended on the endpoint: for clean segmentation and topology metrics, each compared method was paired with FilAdh on the same source images; for count and morphometry, each prediction was paired with its corresponding expert-curated reference. Thus, the analysis used one statistical test throughout, with different prespecified comparison pairs appropriate to each endpoint.

Holm adjustment was applied within each displayed comparison family. The clean FA and SF benchmark families contained 16 tests (four compared methods *×* four metrics) and 12 tests (four compared *×* three metrics), respectively. The morphometry families contained 10 tests in the clean FA table, 15 in the clean SF table, eight in the main FA degradation table, and 12 in the main SF degradation table. Statistical significance was assessed at Holm-adjusted *α* = 0.05; stars denote ^**\***^ *p <* 0.05, ^**\*\***^ *p <* 0.01, and ^**\*\*\***^ *p <* 0.001.

All statistical tests used paired image-level values, with each source image treated as one observation. The estimand was the mean paired difference. A non-significant result was interpreted as failure to detect a difference, not as evidence of equivalence. The main tables present pooled analyses, whereas the supplementary panels show the corresponding comparisons within each dataset. Holm correction was applied within each dataset-specific family for the count and morphometry analyses. Asterisks mark these image-level tests and do not represent tests of individual adhesions, filaments, or density curves. Differences between degradation conditions, classification accuracies, and runtime measurements were summarized descriptively.

## SUPPLEMENTAL INFORMATION

Supplemental information includes twelve figures and two tables containing dataset-stratified benchmark and degradation results, fixed filament-refinement parameters, and synthetic-degradation settings.

## ACKNOWLEDGMENTS

This work was supported in part by the Ministerio de Ciencia, Innovación y Universidades and Agencia Estatal de Investigación (MCIN/AEI/10.13039/501100011033) under Grants PID2023-152631OB-I00 and AIA2025-164165-C41, and in part by the European Regional Development Fund (ERDF), “A way of making Europe.” Dariusz Lachowski was supported by the Consejería de Educación, Ciencia y Universidades, Comunidad de Madrid, under Grant 2023-T1/SAL-GL-29109. This work was also partially supported by the European Union’s Horizon Europe research and innovation programme under grant agreement number 101057970 (AI4Life project). Views and opinions expressed are, however, those of the authors only and do not necessarily reflect those of the European Union or the European Research Council Executive Agency. Neither the European Union nor the granting authority can be held responsible for them. We thank Beatriz Rabadán-Gallego for technical assistance with focal-adhesion and stress-fiber annotation and for reviewing ambiguous image regions.

## AUTHOR CONTRIBUTIONS

Conceptualization, G.R.-P., C.G.-L.-d.-H., D.L., and A.M.-B.; Methodology, G.R.-P. and C.G.-L.-d.-H. with input from D.L. and A.M.-B.; Investigation and annotation, G.R.-P. and M.H.-M.; Formal analysis and visualization, G.R.-P. with input from D.L. and A.M.-B.; Writing – original draft, G.R.-P. and A.M.-B.; Writing – review and editing, all authors; Supervision, D.L. and A.M.-B.

