## Supplemental Figures S1 to S12 and Tables S1 and S2 for "FilAdh: degradation-aware analysis of focal adhesions and actin stress fibers"

### Supplemental information for FilAdh: degradation-aware analysis of focal adhesions and actin stress fibers

#### Supplemental information contents

- Supplemental Figures S1–S3: dataset-stratified focal-adhesion clean-image benchmarks
- Supplemental Figures S4–S6: dataset-stratified stress-fiber clean-image benchmarks
- Supplemental Figures S7–S9: dataset-stratified focal-adhesion degradation analyses
- Supplemental Figures S10–S12: dataset-stratified stress-fiber degradation analyses
- Supplemental Table S1: fixed parameters for orientation-guided filament refinement
- Supplemental Table S2: synthetic-degradation sampling and parameterization

Unless otherwise stated, black dots represent source images and short horizontal bars represent means. The density plots are included to compare the predicted distributions with the expert-curated reference distributions for biologically interpretable measurements, including focal-adhesion area and stress-fiber projected length and apparent width. They show the magnitude and direction of distributional deviation relative to the expert-curated reference; individual adhesions and filaments were not treated as independent experimental units. Asterisks shown in these panels reproduce the corresponding image-level paired comparisons reported in the main analysis and do not represent tests performed on individual components or on the density curves themselves. Confirmatory analyses use the source image as the experimental unit and are described in the main-text Statistical Analysis section. Differences in density-curve shape are otherwise interpreted descriptively.

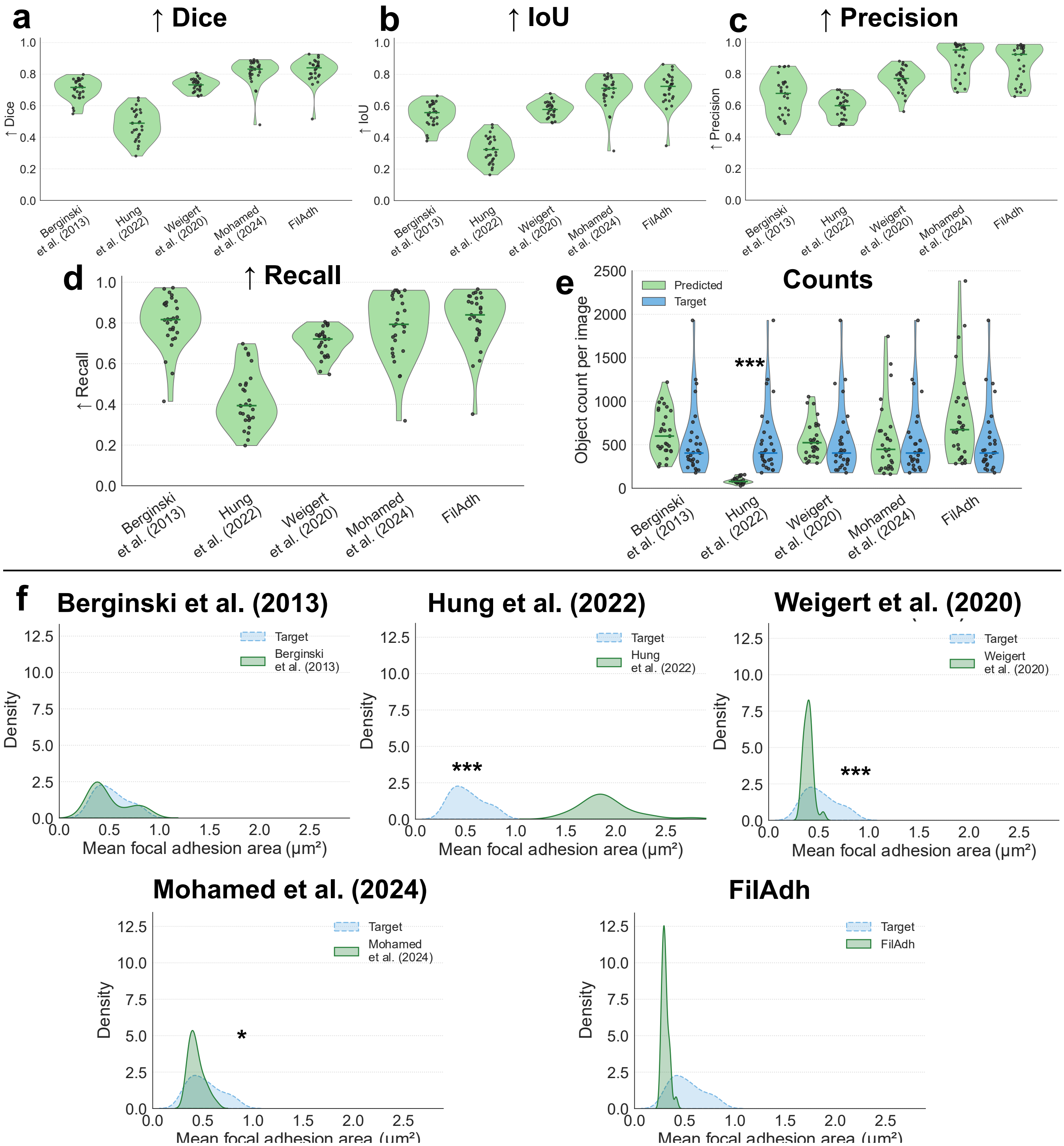


Supplemental Figure S1: **Dataset-stratified focal-adhesion benchmark in HeLa images.** Results are shown for the clean held-out HeLa cohort. **a–d,** Per-image Dice, intersection over union (IoU), precision, and recall for each benchmark method. **e,** Predicted and expert-curated focal-adhesion counts per source image. **f,** Descriptive distributions of per-image mean focal-adhesion area for each prediction method and the unchanged reference. Green denotes predictions, and blue denotes the reference. Black dots represent source images, and short horizontal bars represent means.

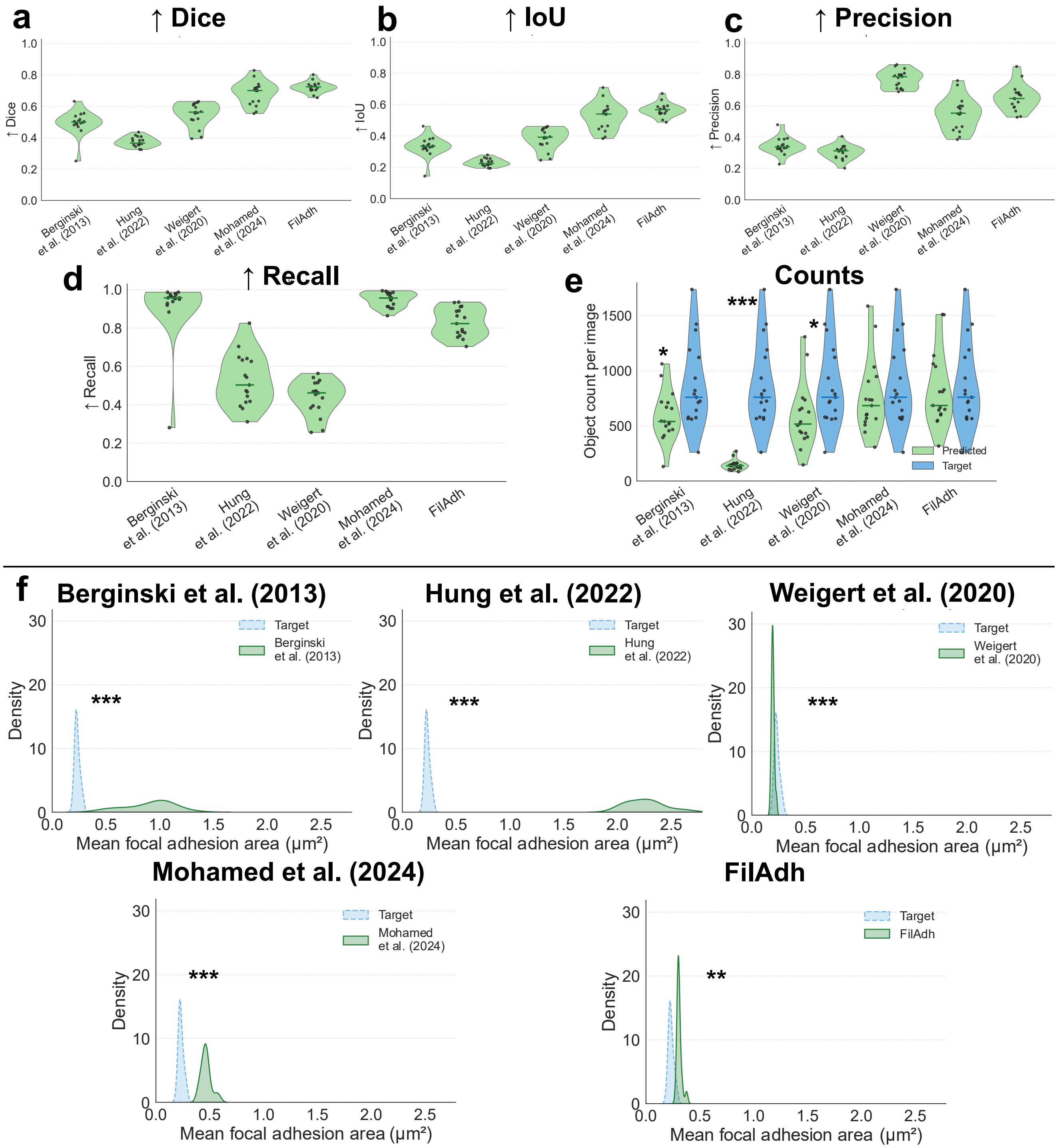


Supplemental Figure S2: **Dataset-stratified focal-adhesion benchmark in SUIT-2 PDAC images.** Results are shown for the clean held-out SUIT-2 cohort. **a–d,** Per-image Dice, intersection over union (IoU), precision, and recall for each benchmark method. **e,** Predicted and expert-curated focal-adhesion counts per source image. **f,** Descriptive distributions of per-image mean focal-adhesion area for each method and the unchanged reference. Green denotes predictions, and blue denotes the reference. Black dots represent source images, and short horizontal bars represent means.

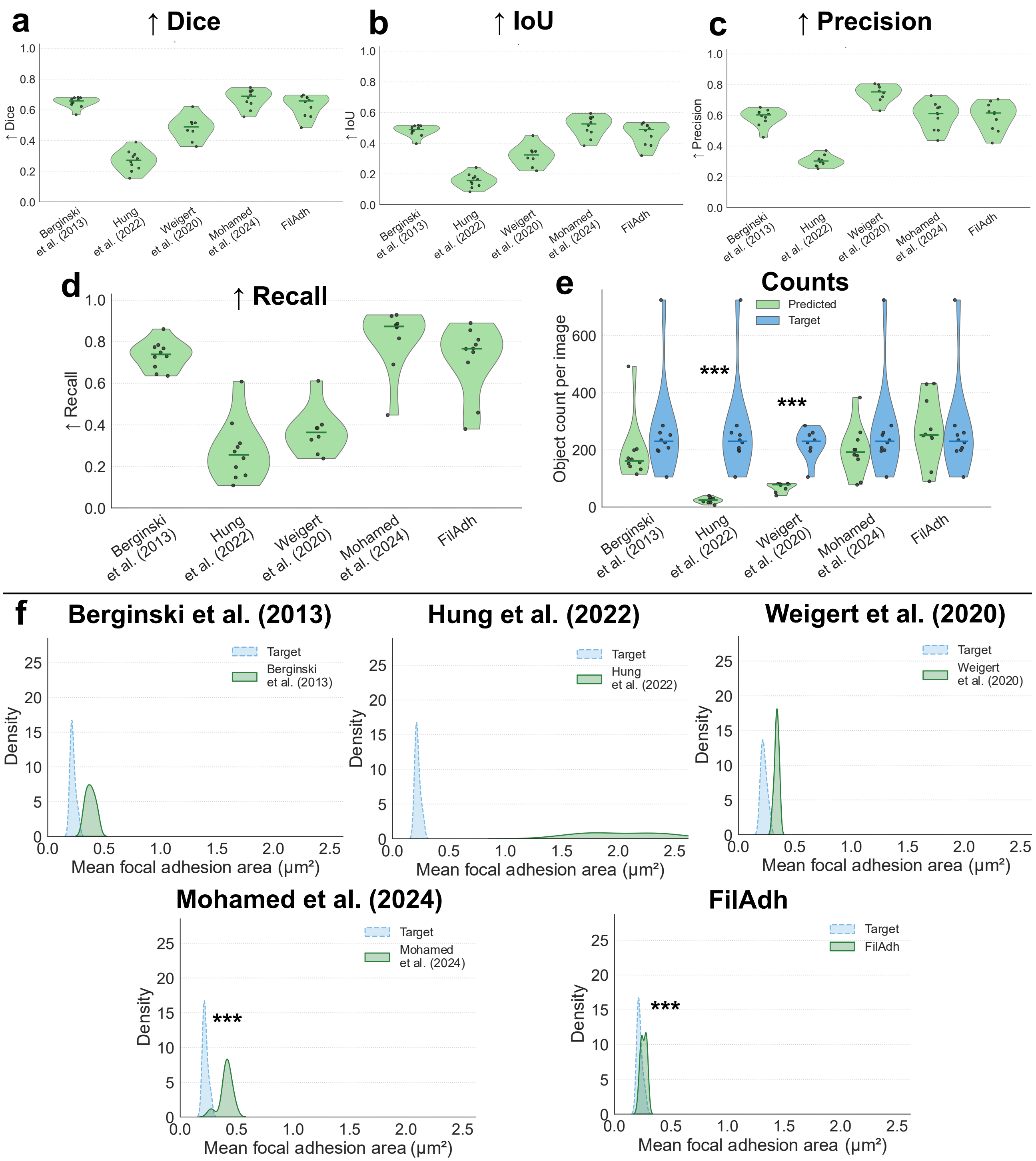


Supplemental Figure S3: **Dataset-stratified focal-adhesion benchmark in MRC-5 images.** Results are shown for the clean held-out MRC-5 cohort. **a–d,** Per-image Dice, intersection over union (IoU), precision, and recall for each benchmark method. **e,** Predicted and expert-curated focal-adhesion counts per source image. **f,** Descriptive distributions of per-image mean focal-adhesion area for each method and the unchanged reference. Green denotes predictions, and blue denotes the reference. Black dots represent source images, and short horizontal bars represent means.

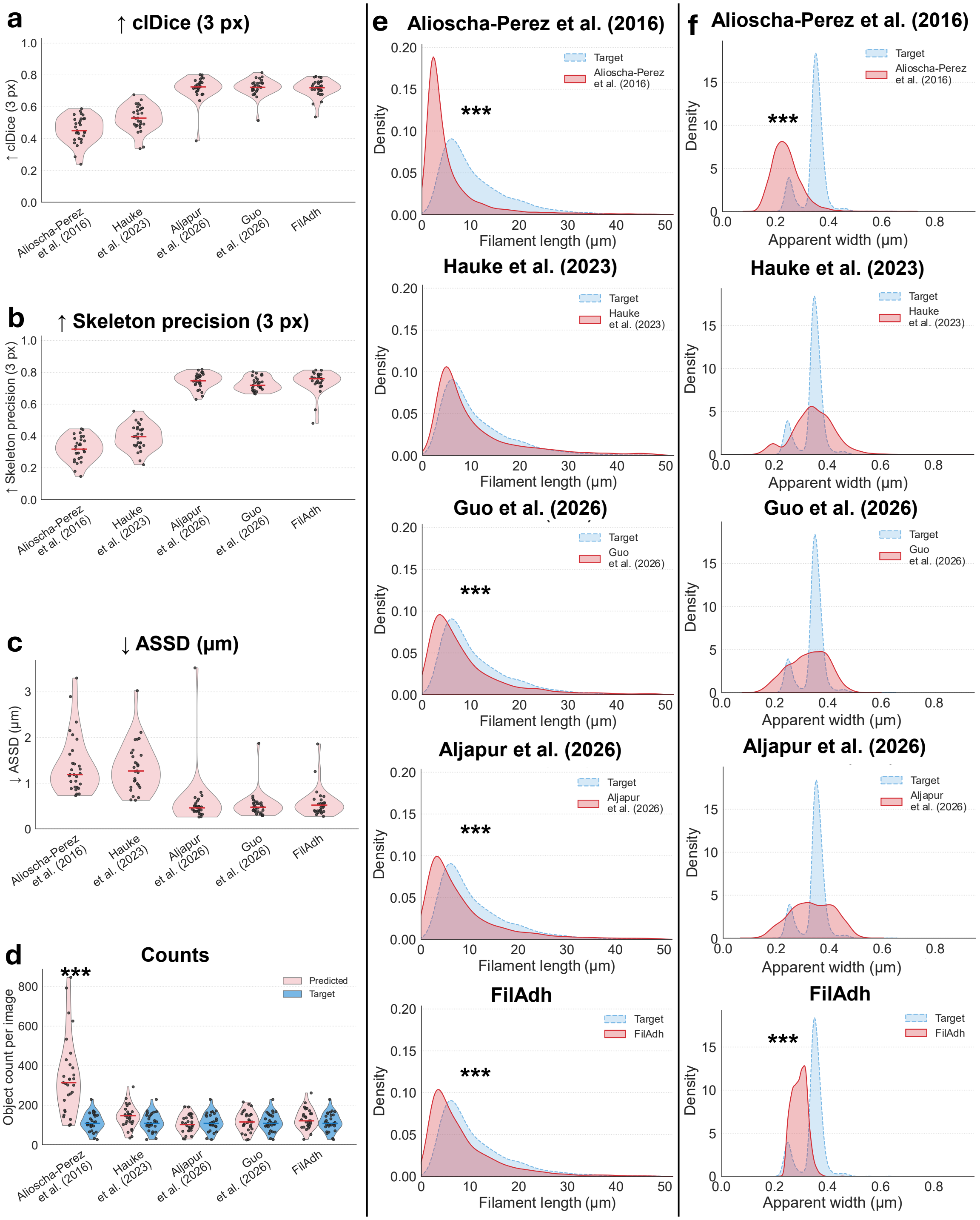


Supplemental Figure S4: **Dataset-stratified stress-fiber benchmark in HeLa images.** Results are shown for the clean held-out HeLa cohort. **a–c,** Per-image 3-pixel-tolerant centerline Dice (clDice), skeleton precision, and average symmetric surface distance (ASSD). **d,** Predicted and expert-curated filament counts per source image. **e,f,** Component-level filament-length and apparent width density plots for each method (red) and the expert-curated reference (blue). These descriptive plots visualize the magnitude and direction of measurement bias relative to the reference and are not component-level inferential comparisons. The distributions illustrate that strong centerline agreement can coexist with count, length, or apparent width bias.

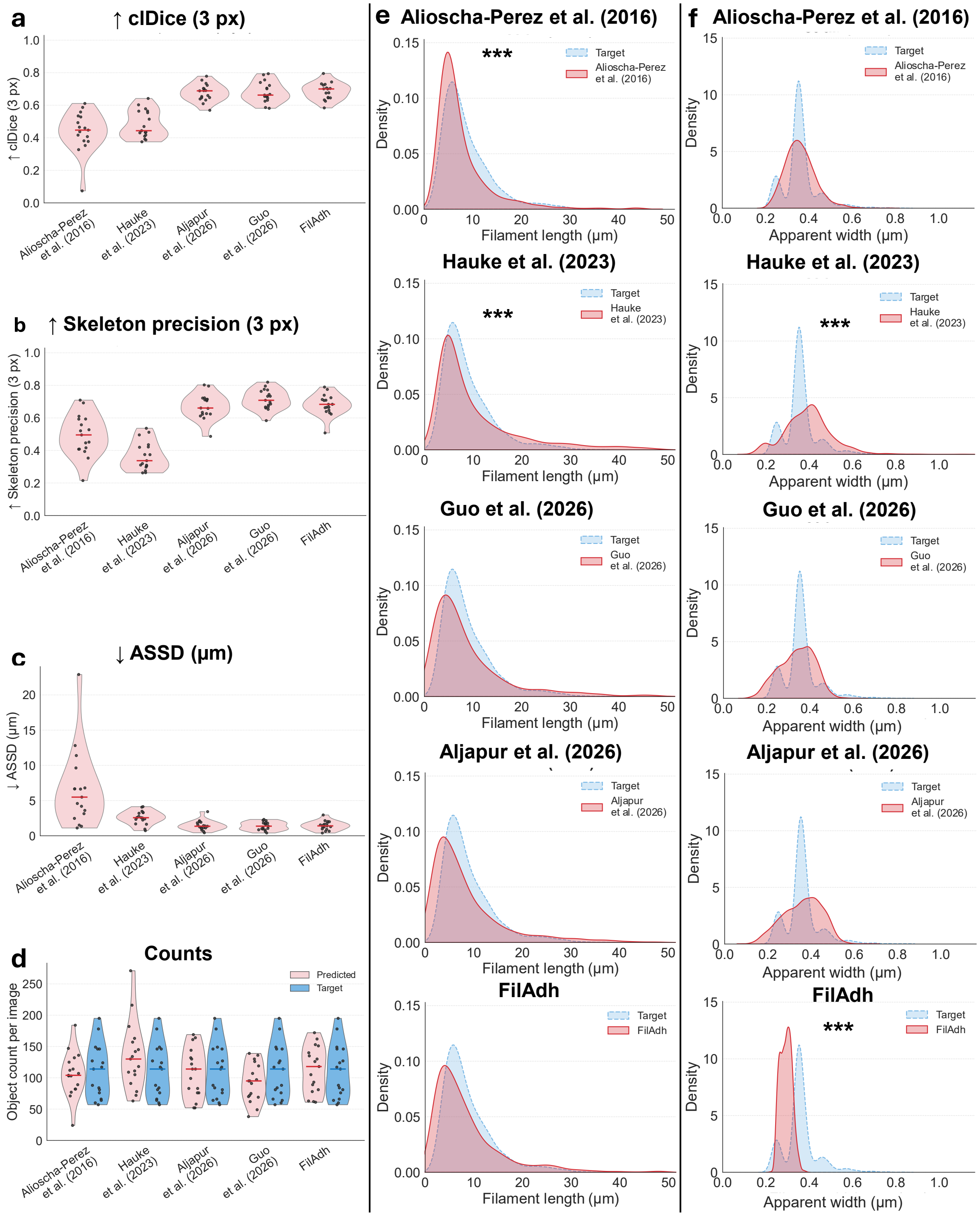


Supplemental Figure S5: **Dataset-stratified stress-fiber benchmark in SUIT-2 PDAC images.** Results are shown for the clean held-out SUIT-2 cohort. **a–c,** Per-image 3-pixel-tolerant centerline Dice (clDice), skeleton precision, and average symmetric surface distance (ASSD) for each benchmark method. **d,** Predicted and expert-curated filament counts per source image. **e,f,** Component-level filament-length and apparent width density plots for each method (red) and the expert-curated reference (blue). These descriptive plots visualize the magnitude and direction of measurement bias relative to the reference and are not component-level inferential comparisons. The distributions illustrate that strong centerline agreement can coexist with count, length, or apparent width bias.

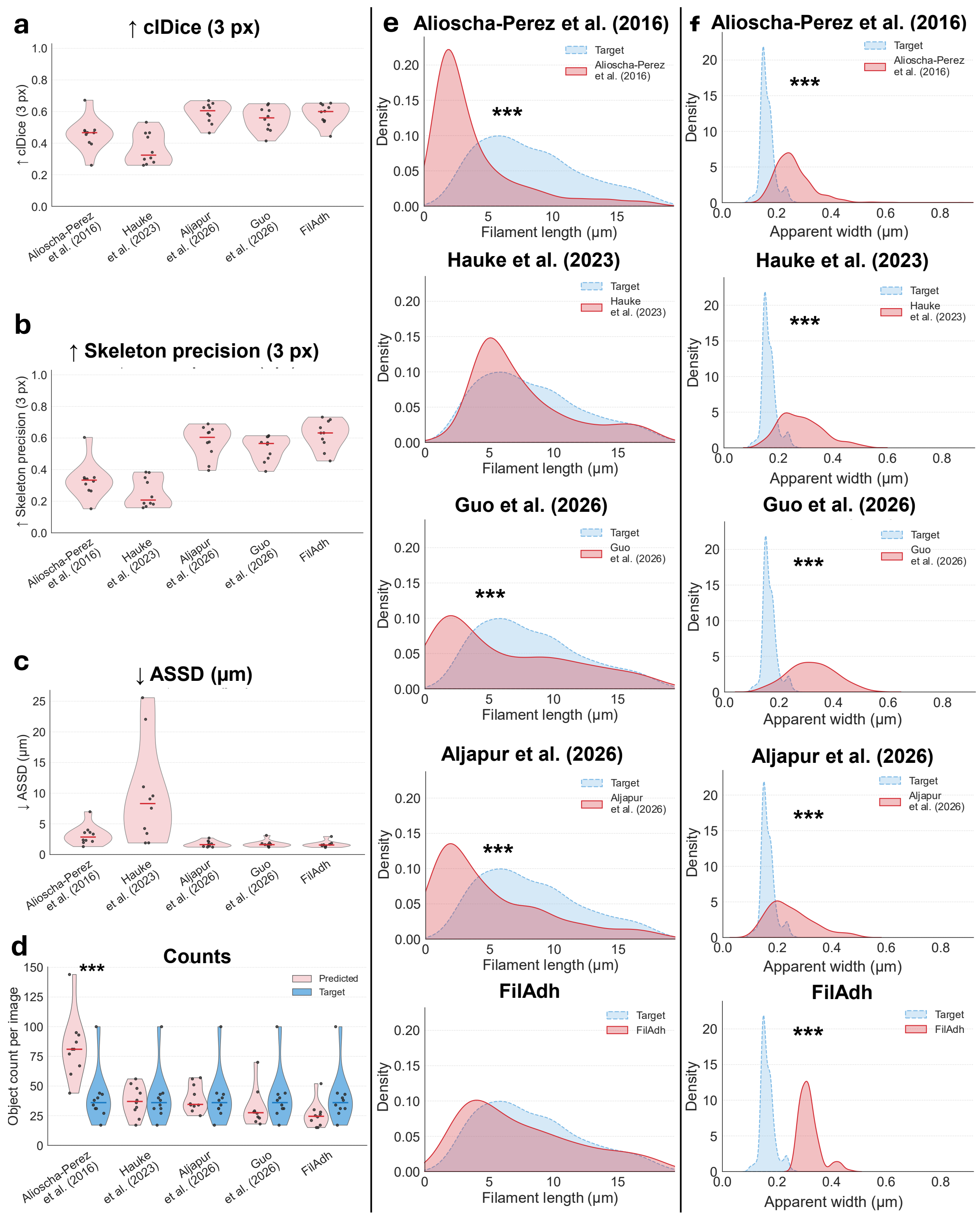


Supplemental Figure S6: **Dataset-stratified stress-fiber benchmark in MRC-5 images.** Results are shown for the clean held-out MRC-5 cohort. **a–c,** Per-image 3-pixel-tolerant centerline Dice (clDice), skeleton precision, and average symmetric surface distance (ASSD) for each benchmark method. **d,** Predicted and expert-curated filament counts per source image. **e,f,** Component-level filament-length and apparent width density plots for each method (red) and the expert-curated reference (blue). These descriptive plots visualize the magnitude and direction of measurement bias relative to the reference and are not component-level inferential comparisons. The distributions illustrate that strong centerline agreement can coexist with count, length, or apparent width bias.

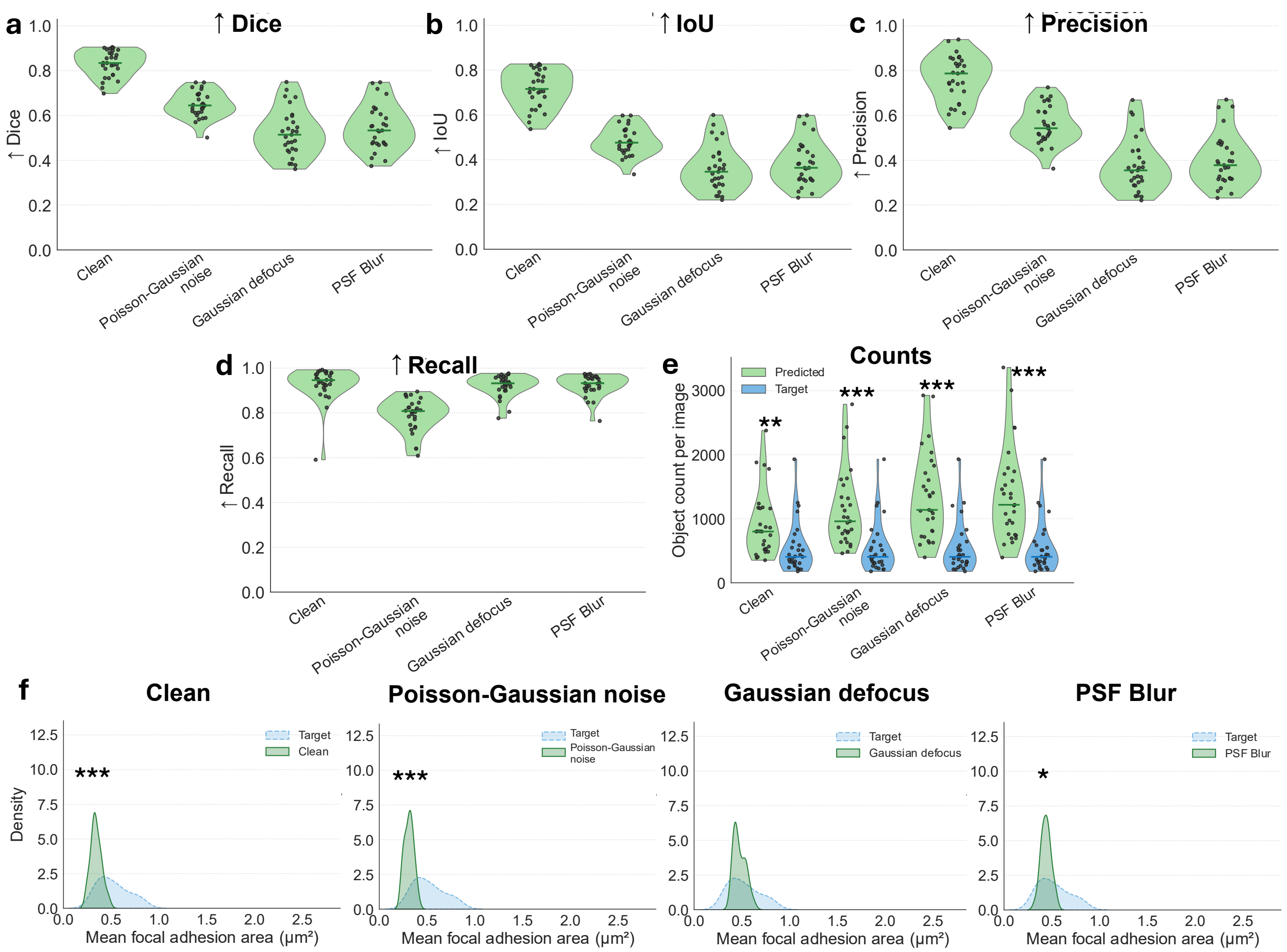


Supplemental Figure S7: **Focal-adhesion degradation analysis in HeLa images.** The degradation-aware FA checkpoint was evaluated on matched clean, Poisson-Gaussian noise, Gaussian defocus, and point-spread-function (PSF) blur versions of each source image. **a–d,** Per-image Dice, intersection over union (IoU), precision, and recall. **e,** Predicted and unchanged expert-curated FA counts per source image. **f,** Descriptive distributions of per-image mean FA area for each condition and the unchanged reference. Green denotes predictions, and blue denotes the reference.

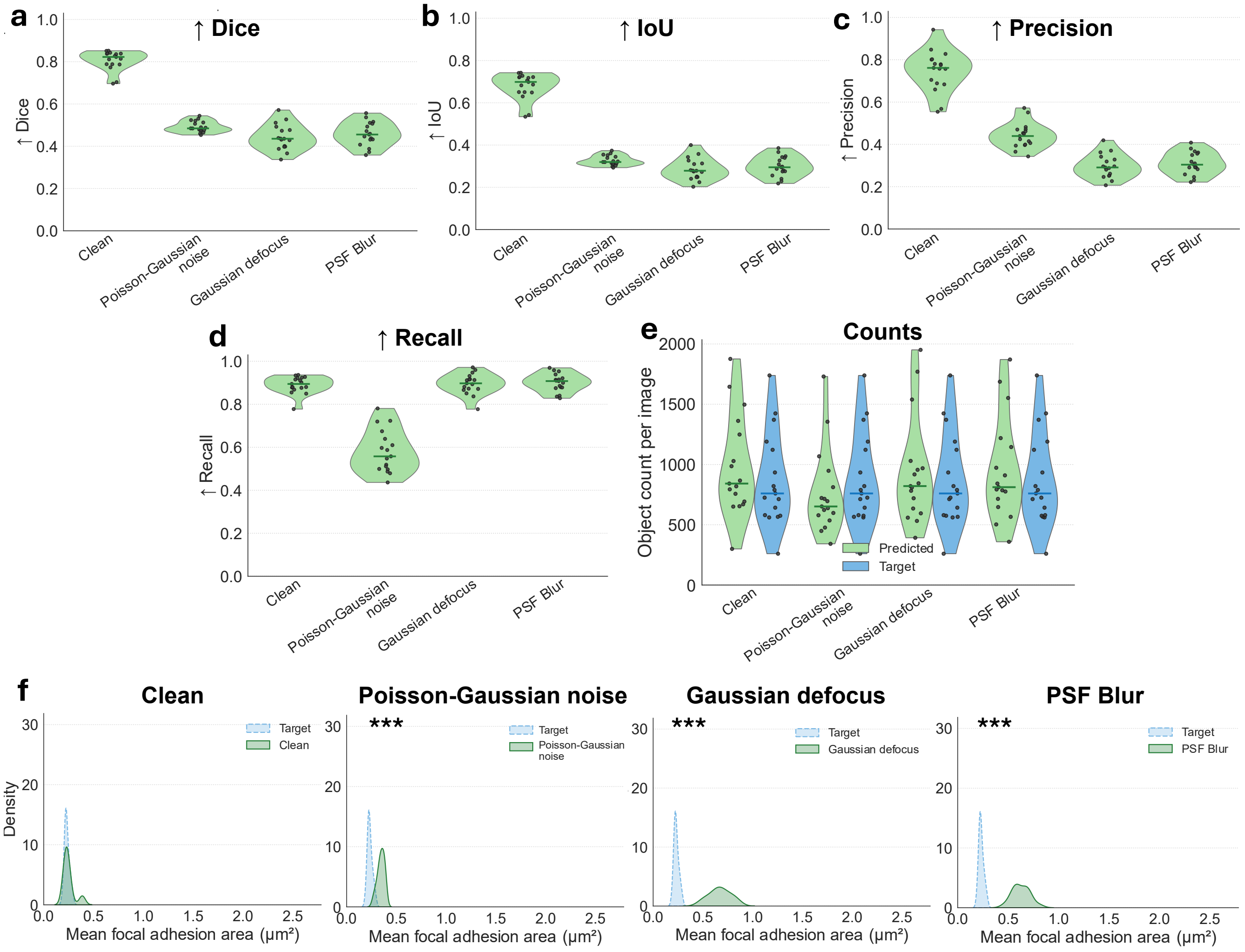


Supplemental Figure S8: **Focal-adhesion degradation analysis in SUIT-2 PDAC images.** The degradation-aware FA checkpoint was evaluated on matched clean, Poisson-Gaussian noise, Gaussian defocus, and point-spread-function (PSF) blur versions of each source image using unchanged reference masks. **a–d,** Per-image Dice, intersection over union (IoU), precision, and recall. **e,** Predicted and reference FA counts per source image. **f,** Descriptive distributions of per-image mean FA area for each condition and the unchanged reference. Green denotes predictions, and blue denotes the reference.

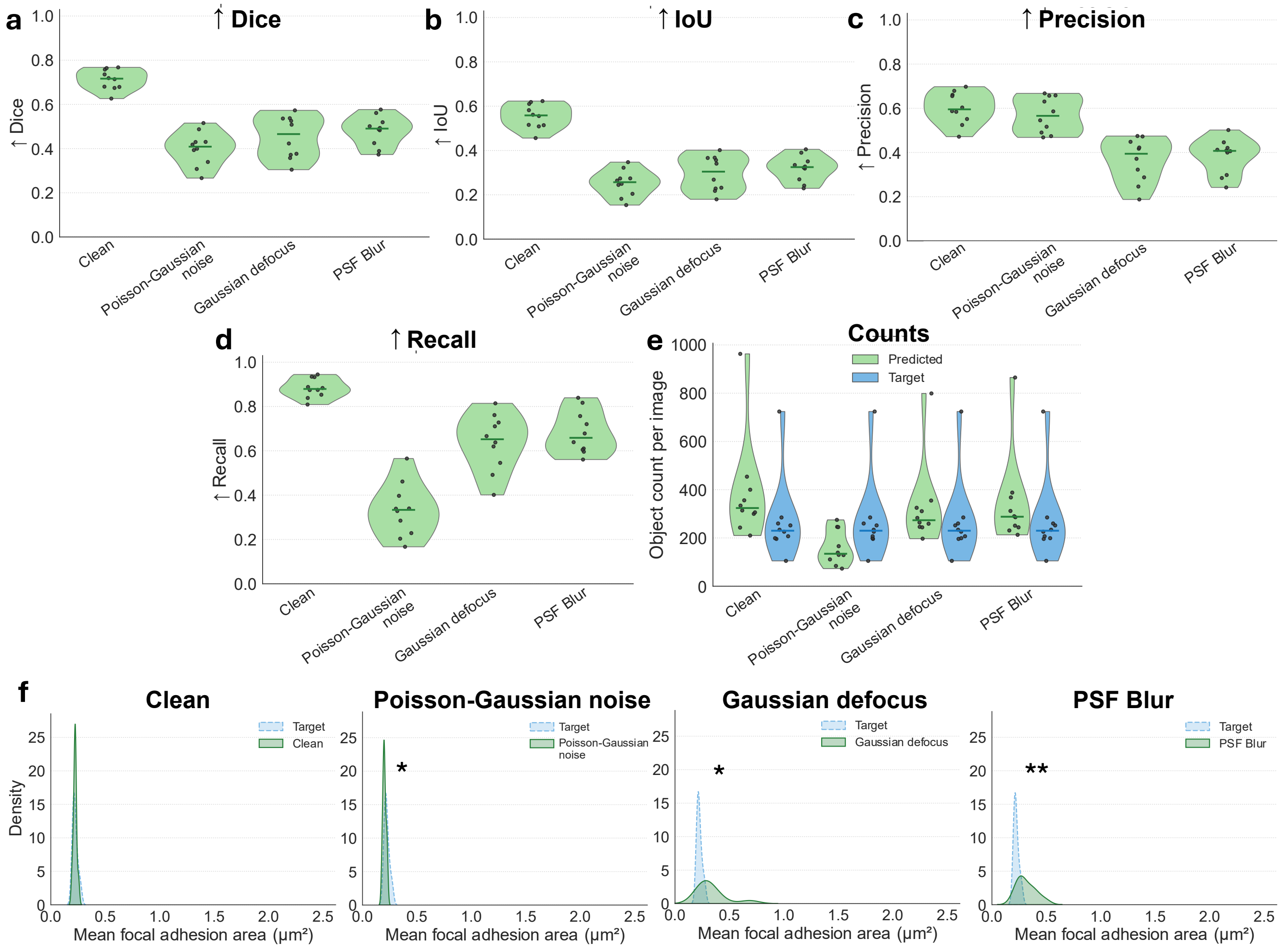


Supplemental Figure S9: **Focal-adhesion degradation analysis in MRC-5 images.** The degradation-aware FA checkpoint was evaluated on matched clean, Poisson-Gaussian noise, Gaussian defocus, and point-spread-function (PSF) blur versions of each source image using unchanged reference masks. **a–d,** Per-image Dice, intersection over union (IoU), precision, and recall. **e,** Predicted and reference FA counts per source image. **f,** Descriptive distributions of per-image mean FA area for each condition and the unchanged reference. Green denotes predictions, and blue denotes the reference.

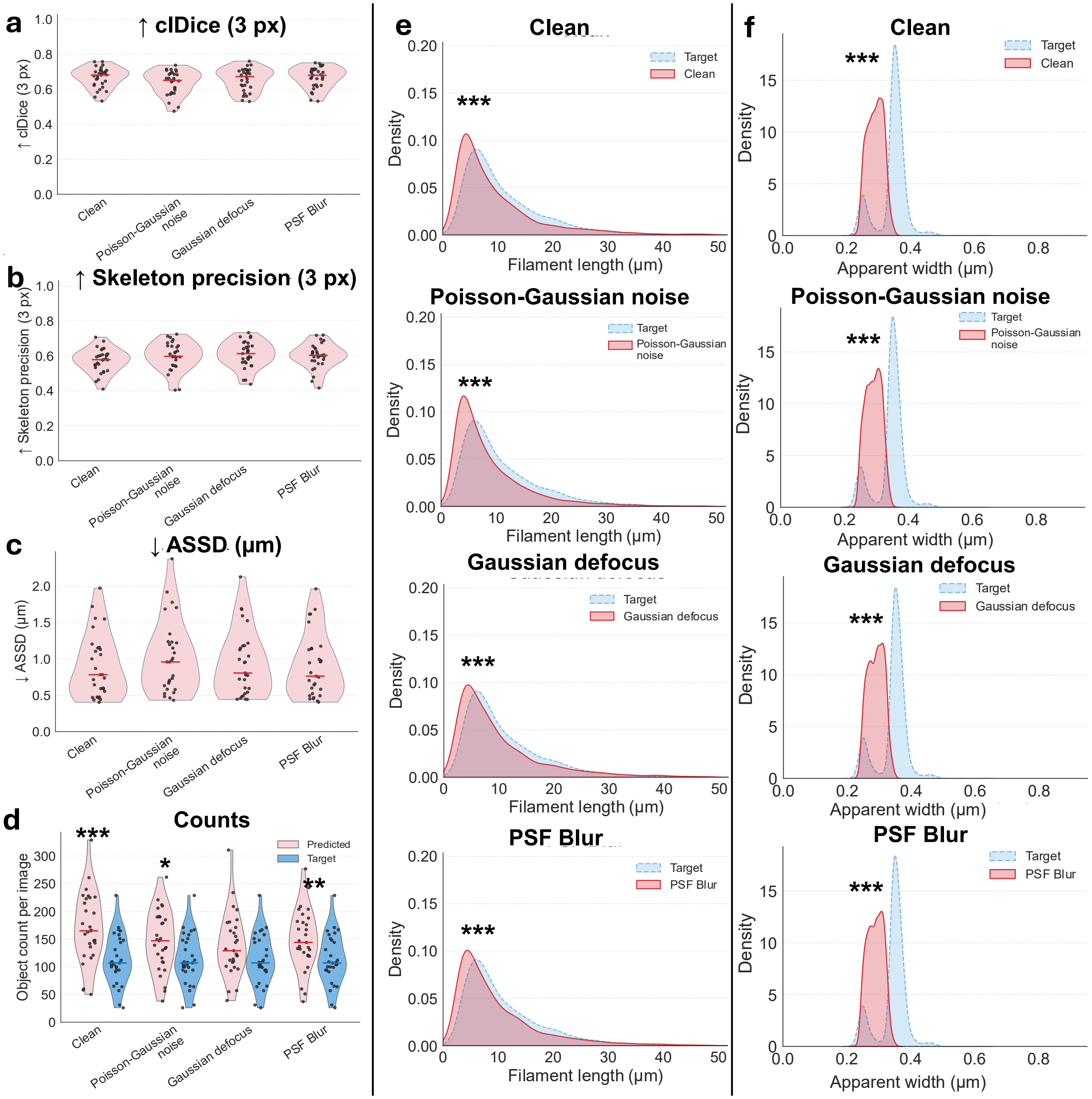


Supplemental Figure S10: **Stress-fiber degradation analysis in HeLa images.** The degradation-aware SF checkpoint was evaluated on matched clean, Poisson-Gaussian noise, Gaussian defocus, and point-spread-function (PSF) blur versions of each source image using unchanged references. **a–c,** Per-image 3-pixel-tolerant centerline Dice (clDice), skeleton precision, and average symmetric surface distance (ASSD). **d,** Predicted and reference filament counts per source image. **e,f,** Descriptive component-level filament-length and apparent width distributions for each condition and the unchanged reference. Pink or red denotes predictions, and blue denotes the reference.

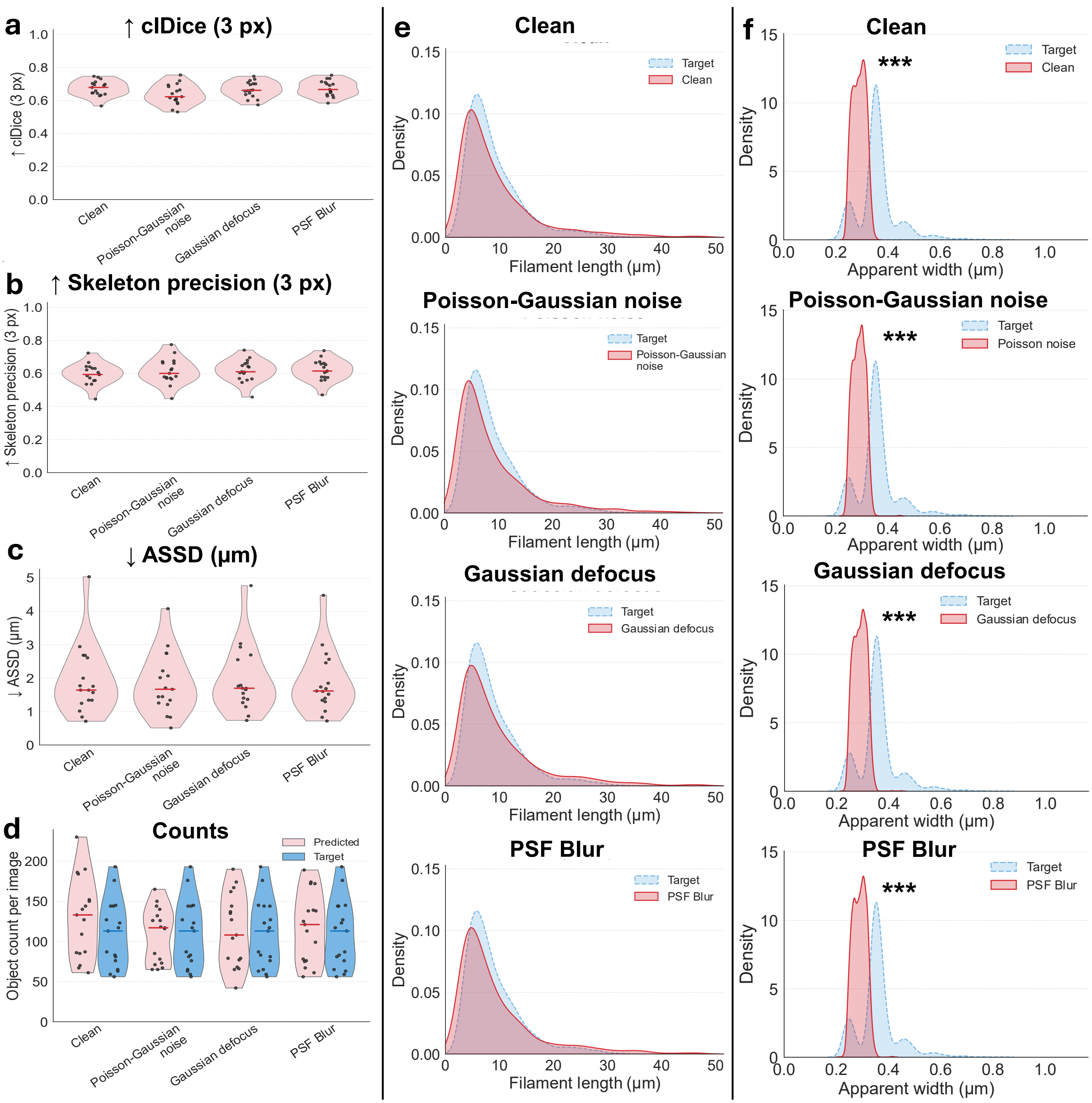


Supplemental Figure S11: **Stress-fiber degradation analysis in SUIT-2 PDAC images.** The degradation-aware SF checkpoint was evaluated on matched clean, Poisson-Gaussian noise, Gaussian defocus, and point-spread-function (PSF) blur versions of each source image using unchanged references. **a–c,** Per-image 3-pixel-tolerant centerline Dice (clDice), skeleton precision, and average symmetric surface distance (ASSD). **d,** Predicted and reference filament counts per source image. **e,f,** Descriptive component-level filament-length and apparent width distributions for each condition and the unchanged reference. Pink or red denotes predictions, and blue denotes the reference.

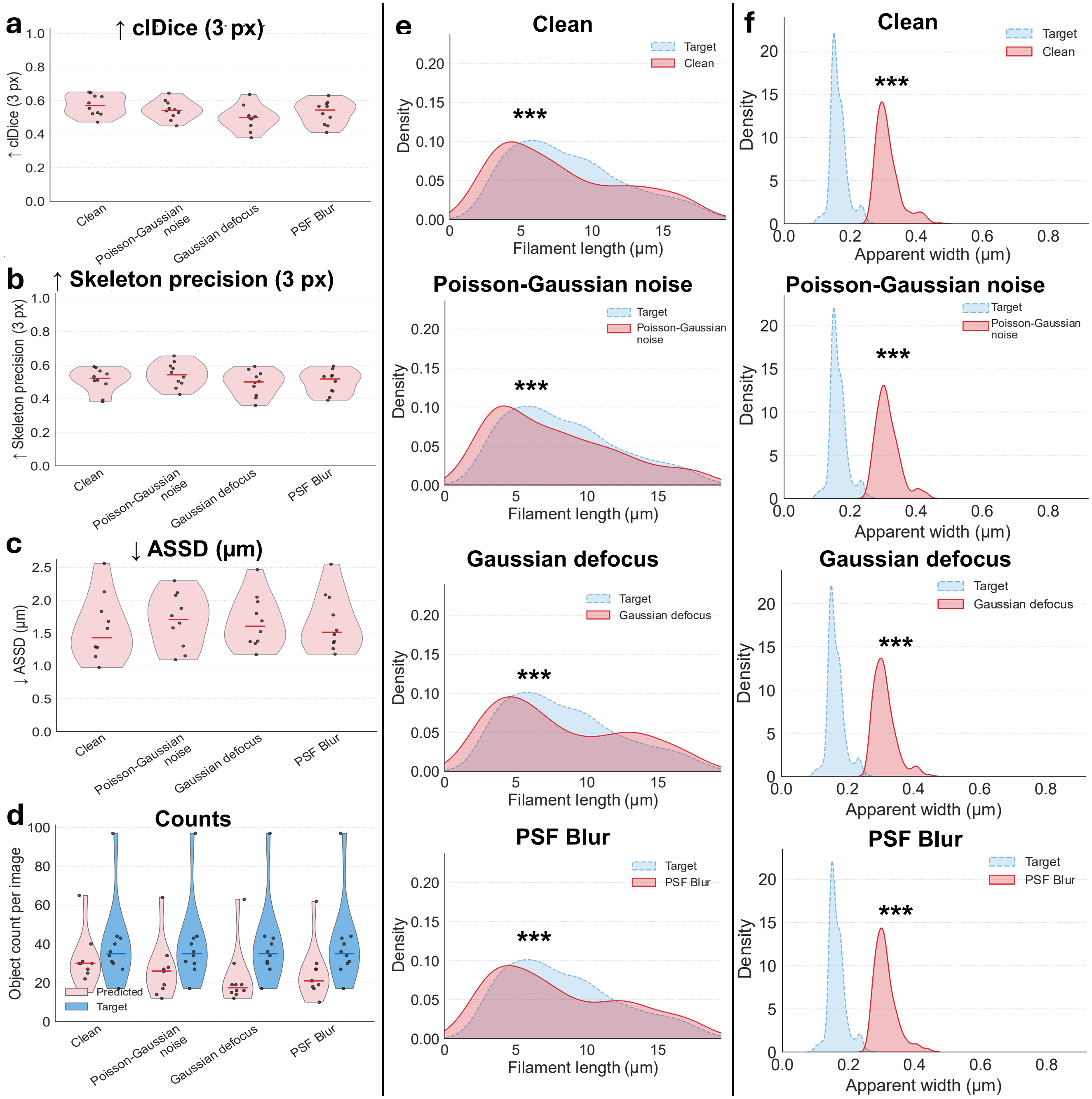


Supplemental Figure S12: **Stress-fiber degradation analysis in MRC-5 images.** The degradation-aware SF checkpoint was evaluated on matched clean, Poisson-Gaussian noise, Gaussian defocus, and point-spread-function (PSF) blur versions of each source image using unchanged references. **a–c,** Per-image 3-pixel-tolerant centerline Dice (clDice), skeleton precision, and average symmetric surface distance (ASSD). **d,** Predicted and reference filament counts per source image. **e,f,** Descriptive component-level filament-length and apparent width distributions for each condition and the unchanged reference. Pink or red denotes predictions, and blue denotes the reference.

#### Supplemental methods tables

Supplemental Table S1: **Fixed parameters for orientation-guided filament refinement.** The parameters were frozen before held-out evaluation and applied unchanged to HeLa, SUIT-2 PDAC, and MRC-5. Pixel-based values therefore represent slightly different physical distances among datasets.

| Parameter | Value | Role in refinement |
|---|---|---|
| Semantic filament threshold | 0.975 | Converts the actin probability map to the binary mask used for instance refinement. |
| Terminal-branch pruning | 10 px | Removes short terminal skeleton branches before junction analysis. |
| Minimum skeleton size | 1.5 $\mu$m | Excludes very short skeleton components. |
| Minimum foreground area | 8 px$^2$ | Excludes very small foreground components. |
| Orientation domain | $[0, \pi)$ | Treats tangent direction as undirected, so orientations separated by $\pi$ are equivalent. |
| Tangent-jump threshold | $40^\circ$ | Marks sharp local orientation changes as candidate path cuts. |
| Angle window | 6 px | Defines the local neighborhood used to evaluate tangent change. |
| Minimum cut spacing | 10 px | Prevents clusters of near-duplicate cuts along the same path. |
| Junction-protection radius | 2 px | Suppresses path cuts close to a detected junction. |
| Minimum junction arms | 2 | Defines the minimum number of sampled arms required for a junction. |
| Arm sampling length | 10 px | Samples local arm direction away from each junction. |
| Maximum bridge distance | 45 px | Limits reconnection across a junction or gap. |
| Maximum geometric-direction difference | $30^\circ$ | Requires opposing arms to be approximately collinear before reconnection. |
| Maximum tangent-orientation difference | $20^\circ$ | Requires the predicted local orientation to support reconnection. |
| Cut radius | 1 px | Defines the radius used to separate incompatible paths. |
| Junction radius | 1 px | Defines the radius used during junction removal. |
| Connectivity | 8-connected | Defines connected components and skeleton neighborhoods. |
| Minimum valid tangent pixels | 2 | Requires sufficient local orientation support for a tangent decision. |
| Initial recovery dilation | Disk radius 1 px | Restores a minimal width around refined skeleton instances. |
| Maximum adaptive dilation iterations | 10 | Recovers instance width within the original semantic mask. |
| Cut-gap policy | Disabled | Allows cut pixels to regrow only when claimed consistently during final constrained dilation. |

Supplemental Table S2: **Synthetic-degradation sampling and parameterization.** The same degradation definitions were used for online degradation-aware training and matched held-out evaluation of the degradation-aware FA and SF checkpoints reported in the main text.

| Component | Parameterization | Purpose |
|---|---|---|
| Severity | $s \sim \mathcal{U}(0.55, 1.35)$ for each online training sample | Controls the strength of the sampled degradation. |
| Training/validation schedule | Clean:PSF:Defocus:Noise $= 1{:}1{:}1{:}3$ | Oversamples low-photon conditions while retaining clean and blurred examples. |
| Held-out testing | One clean, one PSF-blurred, one Gaussian defocused, and one Poisson-Gaussian noise version of each held-out source image for each task | Provides matched condition-level comparisons while retaining the source image as the experimental unit. |
| PSF blur | Airy-like kernel $[2J_1(x)/x]^2$; base radius $r_0 \sim \mathcal{U}(2.0, 4.5)$ px; $r = r_0 \max(0.35, s)$; odd kernel size $\lceil 8r \rceil$ | Provides an Airy-like optical broadening stress test that mimics loss of spatial resolution and small-object boundary contrast. |
| Gaussian defocus | $\sigma_0 \sim \mathcal{U}(0.9, 2.8)$ px; $\sigma = \sigma_0 \max(0.35, s)$ | Approximates defocus and acquisition blur. |
| Poisson-Gaussian noise | Effective peak count $\mathcal{U}(18, 70)/\max(0.40, s)$, followed by Gaussian read noise with standard deviation $0.006 \max(0.25, s)$ | Approximates photon-shot noise together with additive detector read noise. |
| Boundary handling | Reflected convolution boundaries; outputs clipped to $[0, 1]$ | Avoids artificial edge padding and maintains the normalized intensity range. |
